# Homomeric Q/R edited AMPA receptors conduct when desensitized

**DOI:** 10.1101/595009

**Authors:** Ian D. Coombs, David Soto, Thomas P. McGee, Matthew G. Gold, Mark Farrant, Stuart G. Cull-Candy

## Abstract

Desensitization is a canonical property of ligand-gated ion channels, causing progressive current decline in the continued presence of agonist. AMPA-type glutamate receptors, which mediate fast excitatory signaling throughout the brain, exhibit profound desensitization. Recent cryo-EM studies of AMPAR assemblies show their ion channels to be closed in the desensitized state. Here we report the surprising finding that homomeric Q/R edited AMPARs still allow ions to flow when the receptors are desensitized. GluA2(R) expressed alone, or with auxiliary subunits (γ-2, γ-8 or GSG1L), generates large steady-state currents and anomalous current-variance relationships. Using fluctuation analysis, single-channel recording, and kinetic modeling we demonstrate that the steady-state current is mediated predominantly by ‘conducting desensitized’ receptors. When combined with crystallography this unique functional readout of a hitherto silent state enabled us to examine cross-linked cysteine mutants to probe the conformation of the desensitized ligand binding domain of functioning AMPAR complexes within the plasma membrane.

## Introduction

AMPARs mediate fast excitatory signaling in the brain, and a change in their number or function underlies lasting forms of synaptic plasticity^1,2^. At many central synapses the time course of the excitatory postsynaptic current (EPSC) reflects the rapid deactivation of AMPARs following fast neurotransmitter clearance from the cleft^3–5^. AMPAR desensitization, where the channel closes while glutamate remains bound, is also important in shaping transmission, especially during periods of high-frequency synaptic input^6^ or when glutamate clearance is slow^7,8^. In this situation, AMPAR-mediated responses are depressed and AMPARs must recover from desensitization before they can be re-activated^8–10^. Thus, the balance between AMPAR desensitization and recovery influences the amplitude, duration and frequency of neuronal responses^11^.

AMPARs are homo- or heterotetrameric assemblies of the pore-forming subunits GluA1-4. The activation, deactivation and desensitization of the receptor is controlled by ligand binding domains (LBDs) which form a self-contained ‘clamshell’-like structure within each of the four subunits^12,13^. Glutamate binds between the upper (D1) and lower (D2) lobes of these structures. Within the resting receptor, LBDs of adjacent subunits form dimers that are linked ‘back-to-back’ between their D1 domains^12,14^. Following glutamate binding, closure of the LBD clamshell around the agonist causes separation of the D2 domains, applying tension to linkers between the LBDs and the ion channel which opens the gate^12,15–17^. This can be followed by desensitization, which is initiated by rupturing of the D1-D1 interfaces, relieving the tension on the pore linkers imposed by glutamate binding, allowing the channel to close ^14,16,18,19^.

In neurons, AMPARs are intimately associated with numerous classes of auxiliary subunits, which include the transmembrane AMPAR regulatory proteins (TARPs)^20,22^ and germ cell-specific gene 1-like protein (GSG1L)^21^. These auxiliary proteins determine many biophysical and pharmacological properties of AMPARs and influence their desensitization^23,24^. The prototypical TARP γ-2 markedly slows the rate of AMPAR desensitization and accelerates recovery from desensitization^25,26^, while TARP γ-8 and GSG1L slow both the entry into and the recovery from desensitization^21,22,27^. The structures of desensitized complexes, composed of homomeric GluA2 AMPARs with either γ-2 or GSG1L, have recently been determined at ∼8 Å resolution by cryo-electron microscopy (cryo-EM)^16,19^. The desensitized structures displayed a closed pore, a ruptured D1 interface and a modest rearrangement (‘relaxation’) of the LBD dimer with closely apposed D2 lobes^18^. This con-trasts with the more variable LBD structures of GluA2 seen in the absence of auxiliary subunits^28,29^.

Native GluA2 is subject to RNA editing which causes a switch from the genetically encoded glutamine (Q) to an arginine (R) in the selectivity filter; this Q/R editing reduces channel conductance and Ca^2+^ permeability^30–32^. Structural details of the closed, desensitized and activated states of TARP γ-2-associated homomeric GluA2(R) or GSG1L-associated GluA2(Q) have been well characterized^16,17,19,33^. Here we describe striking differences in the functional properties of homomeric GluA2(Q) and GluA2(R) receptors. The edited (R) form displays unusual desensitization behavior and conductance when compared with the unedited (Q) form, or indeed any other AMPAR assembly that we have examined^34–38^. Specifically, we find that GluA2(R) displays a particularly large steady-state current and an anomalous current-variance relationship. When GluA2(R) is expressed with the TARPs (γ-2 or γ-8) or with GSG1L we observe similarly anomalous behavior. We attribute this behavior to pore loop arginines preventing desensitization-mediated channel closure of the GluA2(R) assemblies, giving rise to conducting desensitized receptors. Using functional cysteine cross-linking we exploited this phenomenon to gain insight into the structure of desensitized AMPARs in their native environment.

## Results

### Atypical channel behavior of Q/R edited GluA2

When recording glutamate-evoked currents (10 mM, 100 ms, –60 mV) in outside-out patches excised from HEK293 cells expressing homomeric GluA2 with and without γ-2, γ-8 or GSG1L, we observed unexpected differences in the behavior of the unedited (Q) and edited (R) forms (**Fig. 1**). Compared with that of the Q form, desensitization of the R form was slower – the weighed desensitization time constant (*τ*_w, des_) was greater for all GluA2(R) combinations examined (GluA2 alone +86%, GluA2/γ-2 +26%, GluA2/γ-8 +55%, GluA2/GSG1L +32%) (**Fig. 1a-c**). This slowing of desensitization by Q/R editing was accompanied by a striking increase in fractional steady-state current for GluA2, GluA2/γ-2 and GluA2/γ-8 (*I*_SS_ +470%, +560%, +380%, respectively); by contrast, for GluA2/GSG1L *I*_SS_ was not increased (**Fig. 1a,b,d**).

**Figure 1.**
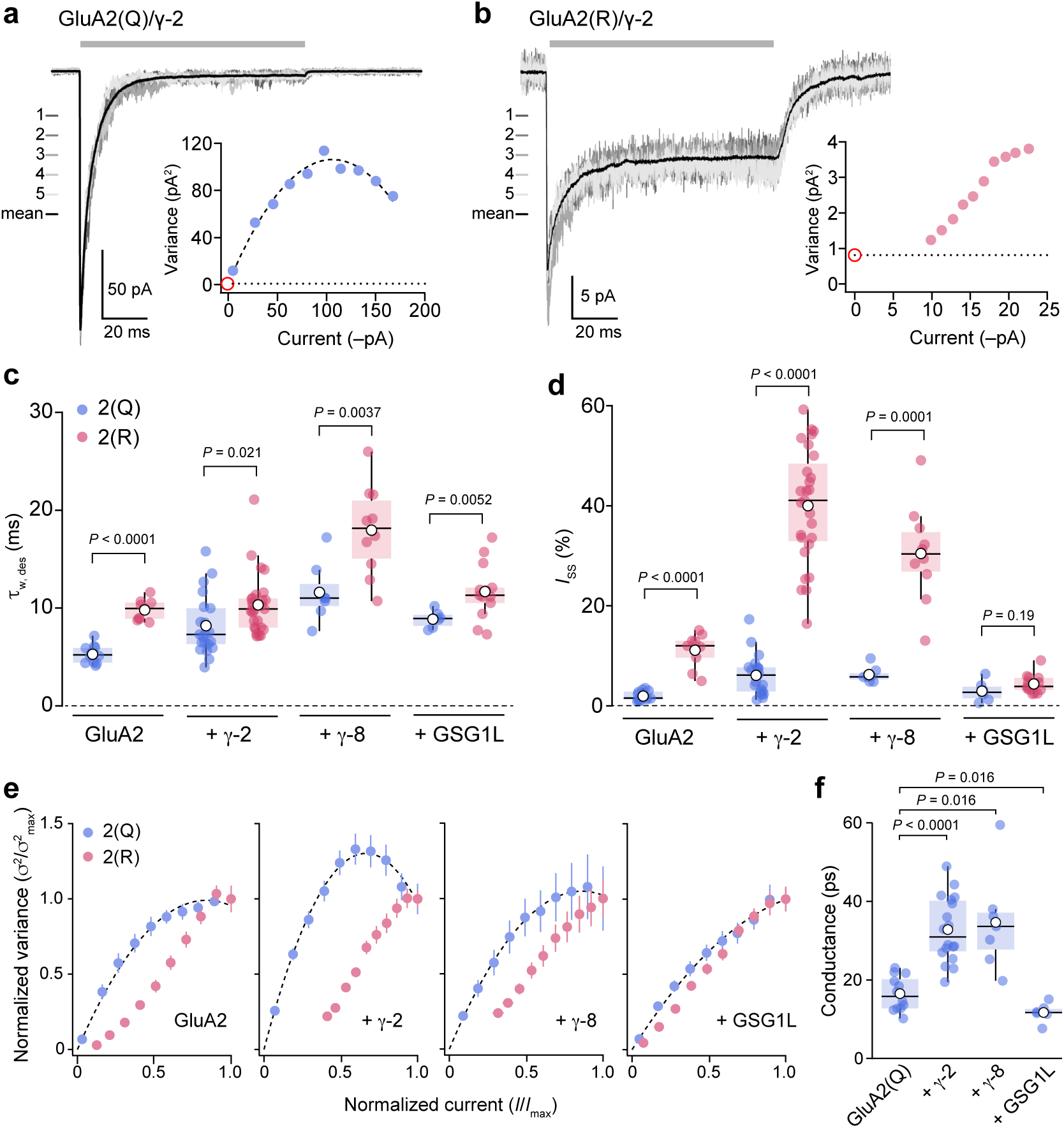
Q/R editing affects the kinetics and variance of GluA2 currents. **a**) Representative outside-out patch responses (10 mM glutamate, 100 ms, –60 mV; gray bar) from a HEK293 cell transfected with GluA2(Q)/γ-2 (average current, black; 5 individual responses, grays). Inset: current-variance relationship (dotted line indicates background variance and red circle indicates expected origin). **b**) As a, but for GluA2(R)/γ-2. Note that the data can not be fitted with a parabolic relationship passing through the origin. **c**) Pooled *τ*_w, des_ data for GluA2 alone (*n* = 12 Q-form and 9 R-form), GluA2/γ-2 (*n* = 21 and 27), GluA2/γ-8 (*n* = 7 and 10), and GluA2/GSG1L (*n* = 6 and 13). Box-and-whisker plots indicate the median (black line), the 25–75th percentiles (box), and the 10–90th percentiles (whiskers); filled circles are data from individual patches and open circles indicate means. Two-way ANOVA indicated: *F* _1,97_ = 111.34, *P* < 0.0001 for Q/R editing; *F* _3,97_ = 32.3, *P* < 0.0001 for auxiliary subunit type; *F* _3,97_ = 2.84, *P* = 0.041 for the interaction of Q/R editing and subunit type. Indicated *P* values are from pairwise Welch *t* -tests. **d**) Pooled data for *I*_SS_. Box-and-whisker plots and *n* numbers as in c. Two-way ANOVA indicated: *F* _1,97_= 129.98, *P* < 0.0001 for Q/R editing; *F* _3,97_ = 58.30, *P* < 0.0001 for auxiliary subunit type; *F* _3,97_ = 58.67, *P* < 0.0001 for the interaction of Q/R editing and subunit type. Indicated *P* values are from pairwise Welch *t* -tests. **e**) Doubly normalized and averaged current-variance relationships (desensitizing current phase only) from GluA2(Q) and GluA2(R) expressed alone (*n* = 12 and 9), with γ-2 (*n* = 19 and 23), with γ-8 (*n* = 7 and 10), or with GSG1L (*n* = 6 and 13). Error bars are sems. All Q-forms can be fitted with parabolic relationships passing through the origin, while R-forms can not. **f**) Pooled NSFA conductance estimates for GluA2(Q) alone, GluA2(Q)/γ-2, GluA2(Q)/γ-8, and GluA2(Q)/GSG1L (*n* = 12, 18, 7 and 6, respectively). Box-and-whisker plots as in c. Indicated *P* values are from pairwise Welch *t* -tests.

In order to determine the underlying weighted mean single-channel conductance for the Q and R forms of the receptors we used non-stationary fluctuation analysis (NSFA) (**Fig. 1a,b,e**). Such analysis typically produces current-variance relationships that can be fitted by a parabolic function which extrapolates to the origin^36,37^. All GluA2(Q) combinations yielded plots with these features. The estimated weighted-mean single-channel conductance (16.5 ± 1.3 pS, *n* = 12) was increased by co-expression with γ-2 or γ-8 (32.8 ± 2.0 pS and 34.6 ± 4.8 pS; *n* = 18 and 7, respectively), but reduced by co-expression with GSG1L (11.7 ± 1.0 pS, *n* = 6) (**Fig. 1f**), consistent with previous reports^37,38^. By contrast, NSFA of GluA2(R) receptor combinations produced anomalous current-variance relationships that were right-shifted (**Fig. 1b,e**), precluding conventional interpretation. This shift, which was apparent for GluA2(R) alone, was accentuated by expression with TARPs γ-2 or γ-8, but was reduced by co-expression of GSG1L (**Fig. 1e**). We have not previously seen such current-variance relationships with other AM-PAR complexes^34–38^.

To determine the basis of the NSFA anomalies, we focused on GluA2(R)/γ-2, which displayed the most robust expression, the greatest increase in steady-state current and the most right-shifted current-variance relationship. First, we observed a near identical current-variance relationship for the tandem construct GluA2(R)_γ-2 (**Supplementary Fig. 1a**). This suggests that the behavior of co-expressed GluA2(R) and γ-2 did not simply reflect the presence of AMPARs with different TARP stoichiometries, and thus heterogeneous channel properties^39^. Second, shifted relationships were also seen with GluA2(R)/γ-2 at +60 mV (**Supplementary Fig. 1b,c**). As channels would be passing Cs^+^ rather than Na^+^ in this condition, this argues that the phenomenon is independent of both voltage and permeating ion. Third, we obtained anomalous current-variance relationships with the edited form of GluA4 (GluA4(R)/γ-2) (**Supplementary Fig. 1d**), indicating that the behavior is not confined to GluA2(R) receptors. Fourth, current-variance relationships derived from deactivation of GluA2(R)/γ-2 (following 1 or 100 ms glutamate exposure; **Supplementary Fig. 2a-e**) and from GluA2(R)_γ-2 or GluA4(R)/γ-2 (following 1 ms glutamate exposure; **Supplementary Fig. 2f,g**) also displayed non-parabolic features. Taken together, our results reveal that the behavior of homomeric Q/R-edited AMPARs deviates substantially from that expected.

**Figure 2.**
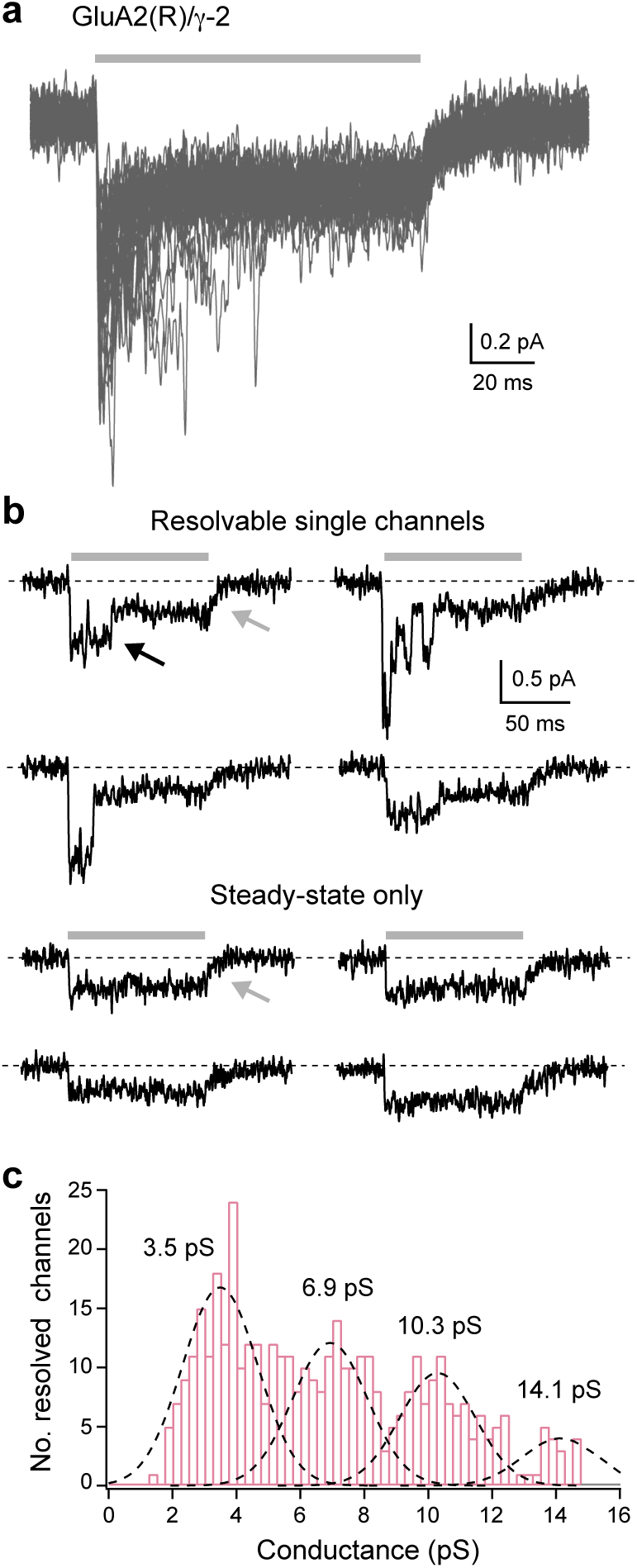
GluA2(R)/γ-2 single-channel recordings. **a**) GluA2(R)/γ-2 currents from an outside-out patch containing few channels (–60 mV). Forty consecutive applications of 10 mM glutamate (gray bar) are overlaid. **b**) Individual responses exhibiting discrete channel openings riding on a persistent steady-state current (upper sweeps) or exhibiting only a persistent steady-state component (lower sweeps). Note the decay of the steady-state currents on glutamate removal (gray arrows) is much slower than the closure of resolved channels (black arrow). **c**) Histogram of channel conductance for 392 discernible openings from 6 patches. The histogram is fit with the sum of four Gaussian curves (dashed lines) with a common standard deviation (1.2 pS), revealing four peaks (3.5, 6.9, 10.3 and 14.1 pS).

### GluA2(R)/γ-2 receptors display two distinct types of channel opening

Stationary fluctuation analysis of GluA2(R) currents in the absence of TARPs has previously yielded an estimated conductance of ∼300 fS^32^. While single-channel openings of this magnitude would be too small to resolve directly, in some of our GluA2(R)/γ-2 patches containing small numbers of channels, we were able to observe discrete single-channel openings that were in the picosiemens range (**Fig. 2a,b**). A histogram of channel amplitudes (pooled from six patches) revealed that openings to a conductance level of 3.5 pS were the most prevalent (**Fig. 2c**). The additional peaks above 3.5 pS could reflect either the presence of multiple conductance states, as reported for unedited AMPAR combinations ^26,32,40–42^, or multiple concurrent events.

The records with resolved openings showed several unusual features. First, despite the steady-state current being relatively large (∼40% of the peak current; **Fig. 1d**), the majority of resolvable openings were present at the onset of the glutamate application (**Fig. 2a,b**). Second, while the amplitudes of resolved openings were equivalent to (or larger than) the steady-state current, the recordings contained no sojourns to the baseline; the resolvable openings thus appeared to ‘ride’ on a low-noise background current (**Fig. 2a,b**). Third, throughout the recordings, occasional ‘failures’ were observed in which the current onset showed no discernible picosiemens openings. Nonetheless, these responses still exhibited the low-noise steady-state current (**Fig. 2b**). Fourth, unlike the rapid closure of the resolved picosiemens channel openings, the steady-state current decay upon agonist removal was slow (and roughly exponential), as might be expected if it reflected the closure of a large number of lower conductance openings (**Fig. 2b**). These results suggest that GluA2(R)/γ-2 receptors are capable of generating two distinct types of channel opening – ‘conventional’ AMPAR openings (with conductances in the picosiemens range) which form the initial phase of the macroscopic response, along with openings of much lower conductance, that form the steady-state current. The pattern of channel behavior observed – predominantly large openings occurring at the onset of the response and predominantly small openings at steady-state – would give rise to a steady-state current with relatively low variance, consistent with the right-shifted current-variance relationships (**Fig. 1e**).

GluA2(R) receptors in the absence of auxiliary subunits exhibit detectable chloride permeability (*P*_Cl_/*P*_Cs_ estimated as 0.14; Ref. 43). We asked whether the different classes of channel openings seen with GluA2(R)/γ-2 could have different relative chloride permeabilities. To address this, we applied 100 ms pulses of glutamate and measured the reversal potential of the peak and steady-state currents (comprising mostly large and small openings respectively) in external solutions containing high (145 mM) or low (35 mM) CsCl^43^ (**Supplementary Fig. 3**). The expected shift in reversal potential following a switch between these conditions is –30.4 mV for a Cs^+^-selective (Cl^−^-impermeable) channel and +30.1 mV for a purely Cl^−^-selective channel. We recorded shifts in reversal potential of –36.0 ± 1.4 mV and – 33.7 ± 2.1 mV for the peak and steady-state currents, respectively (*n* = 7; **Supplementary Fig. 3**), suggesting that the different classes of channel opening do not differ in their relative chloride permeability. Indeed, this result suggests that in the presence of γ-2, GluA2(R) receptors mediate negligible Cl^−^ flux.

**Figure 3.**
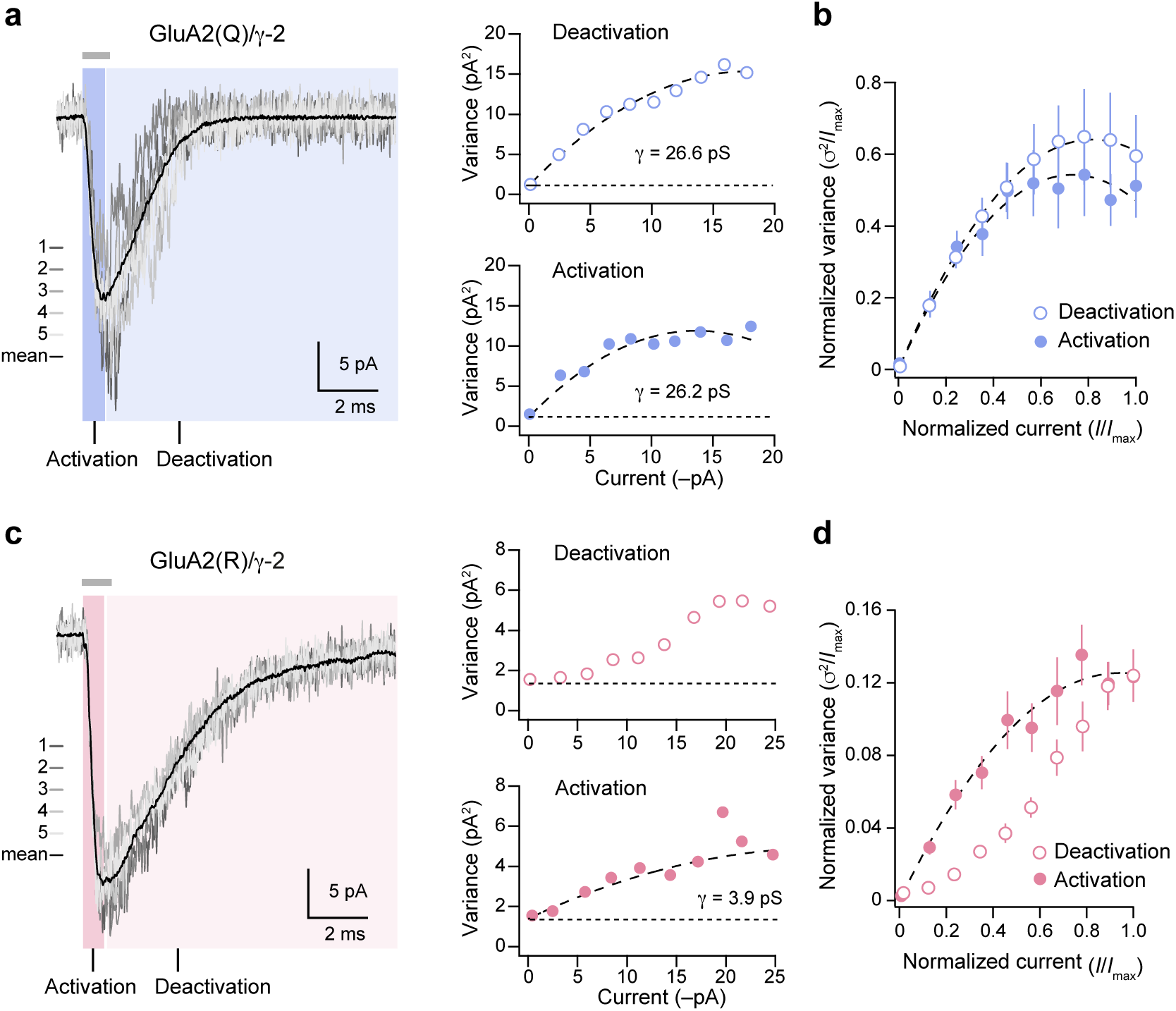
Estimated weighted mean conductance from NSFA of GluA2(R)/γ-2 activation. **a**) Representative GluA2(Q)/γ-2 responses to 1 ms (gray bar) glutamate application (gray traces) with superimposed average (black trace). NSFA was performed on the activation phase (blue highlight; filled blue circles) and deactivation phase (light blue highlight; open blue circles) of the same records, yielding similar estimates of weighted mean conductance. **b**) Pooled current-variance plots for the activation and deactivation of GluA2(Q)/γ-2 currents (*n* = 5). Error bars indicate sem. **c**) Representative GluA2(R)/γ-2 responses to 1 ms glutamate application (as in a). NSFA was performed on the activation phase (pink highlight; filled pink circles) and deactivation phase (light pink highlight; open pink circles) of the same records. Current-variance relationship is non-parabolic for deactivation but parabolic for activation. **d**) Pooled current-variance plots for the activation and deactivation of GluA2(R)/γ-2 currents (*n* = 8). Error bars indicate sem.

### Low-noise steady-state current arises from conducting desensitized channels

NSFA for both desensitization and deactivation gave current-variance relationships that were not amenable to conventional interpretation. We thus sought to determine whether NSFA of the rising phase of the current (AMPAR activation) could accurately report weighted-mean single-channel conductance and, if so, whether this approach might be applicable to GluA2(R)/γ-2. We first confirmed that NSFA of the fast-rising AMPAR activation phase would allow us to estimate accurately the single-channel conductance of unedited GluA2(Q)/γ-2. This yielded a weighted mean conductance of 24.7 ± 4.3 pS, not different from that obtained from the analysis of deactivating current (26.4 ± 3.1 pS, *n* = 5; **Fig. 3a,b**). For GluA2(R)/γ- 2 activation (unlike the deactivation phase of the same records) we obtained conventional parabolic current-variance relationships yielding a weighted mean conductance of 3.8 ± 0.5 pS (*n* = 10; **Fig. 3c,d**), similar to the most prevalent conductance seen in our single-channel analysis (**Fig. 2c**).

As conventional current-variance relationships could be produced from currents recorded during GluA2(R)/γ-2 activation but not desensitization (nor indeed from deactivation records, during which there is a degree of desensitization) we speculated that desensitization itself may provide the key to our unexpected results. Specifically, we considered whether desensitization might not fully close the ion channel, such that the GluA2(R)/γ-2 receptors could adopt a conducting desensitized state, giving rise to the large steady-state current and the anomalous current-variance relationships. If this were the case, and the shift from large resolvable channel openings to smaller openings was linked to the process of desensitization, a decline in GluA2(R)/γ-2 single-channel conductance (and therefore macroscopic current) would not be expected if desensitization was blocked.

In the presence of cyclothiazide, which inhibits desensitization by stabilizing the upper LBD dimer interface^14^, we found that GluA2(R)/γ-2 macroscopic currents (10 mM glutamate, 1 s) did not decay (**Fig. 4a**). Likewise, if the low-noise steady-state current of GluA2(R)/γ-2 arose from conducting desensitized channels, then we would expect the steady-state current to remain when desensitization was enhanced. To test this, we used the point mutation S754D. This weakens the upper LBD dimer interface, accelerating desensitization and reducing steady-state currents of GluA2(Q)^14^. For both GluA2(Q)/γ-2 and GluA2(R)/γ-2 the S754D mutation produced a near 20-fold acceleration of desensitization (**Fig. 4b,c**) and a greater than 2-fold slowing of recovery from desensitization (**Fig. 4d**). As anticipated, GluA2(Q) S754D/γ-2 produced a negligible steady-state current (**Fig. 4b,e**). In marked contrast, GluA2(R) S754D/γ-2 exhibited an appreciable steady-state current (**Fig. 4b,e**). The fact that this exists under conditions strongly favoring desensitization is consistent with the idea that desensitized channels can conduct.

**Figure 4.**
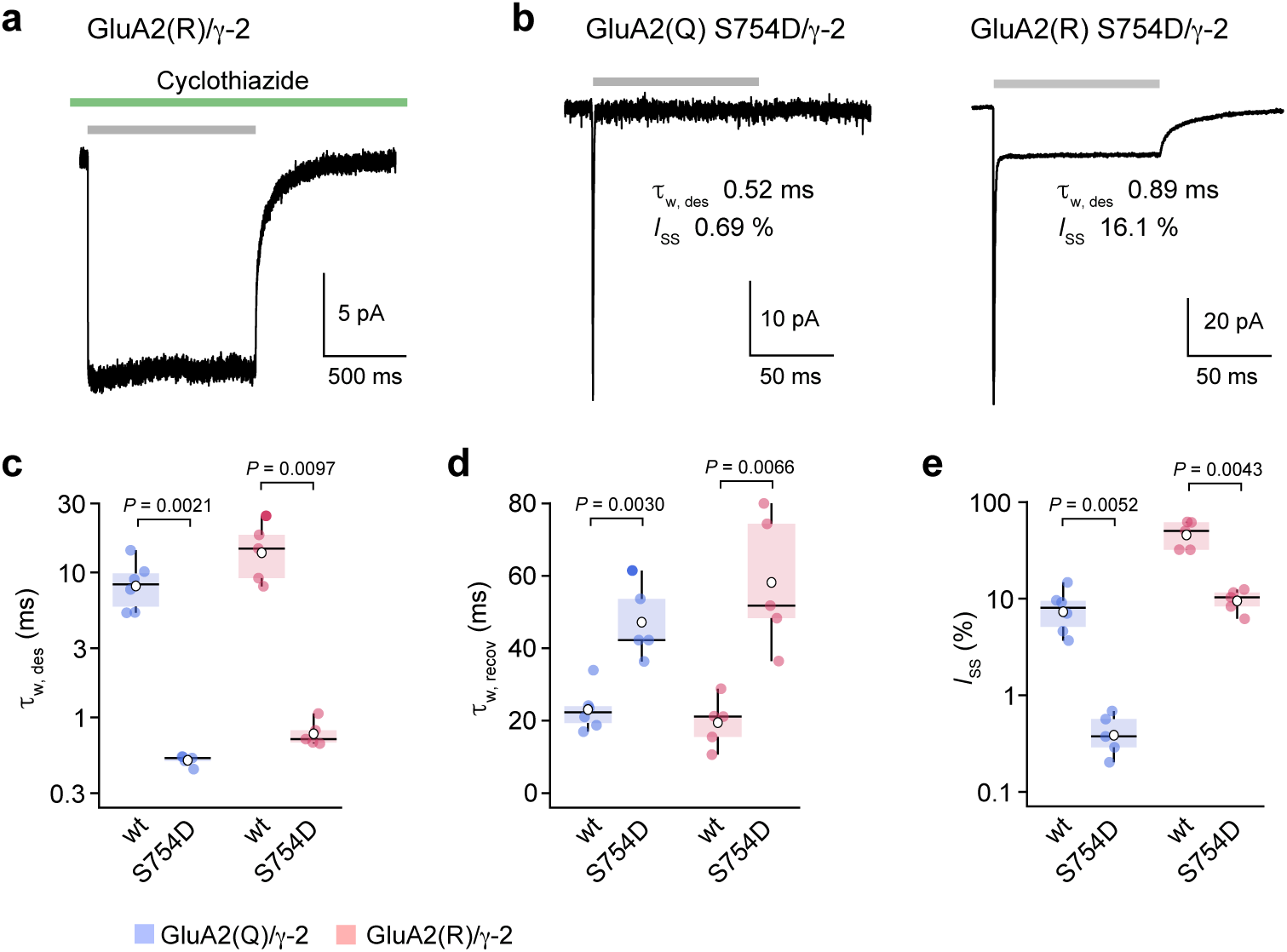
Large steady-state GluA2(R)/γ-2 currents are observed even in conditions favoring desensitization. **a**) Representative GluA2(R)/γ-2 current (–60 mV) evoked by 10 mM glutamate (gray bar) in the presence of 50 μM cyclothiazide (green bar). Note the minimal current decay, as desensitization is inhibited (for pooled data *I*_SS_/*I*_peak_ = 93.4 ± 1.6%, *n* = 6). **b**) Representative glutamate-evoked currents from Q and R forms of GluA2 S754D/γ-2. Both forms exhibit very fast desensitization, but the R form has an appreciable steady-state current. **c**) Pooled data showing desensitization kinetics (τ_w, des_) for wild-type (wt; *n* = 6 and 5) and mutant (S754D; *n* = 5 and 5) forms of GluA2(Q)/γ-2 and GluA2(R)/γ-2. Box-and-whisker plots as in Fig. 1c. Two-way ANOVA indicated: *F* _1, 17_ = 10.56, *P* = 0.0047 for Q/R editing; *F* _1, 17_ = 43.19, *P* < 0.0001 for the mutation; *F* _1, 17_ = 2.63, *P* = 0.12 for the interaction between Q/R editing and the mutation. **d**) Pooled data (as in c) for recovery kinetics (τ_w, recov_). Two-way ANOVA indicated: *F* _1, 17_ = 0.13, *P* = 0.72) for Q/R editing; *F* _1, 17_ = 31.67, *P* < 0.0001 for the mutation; *F* _1, 17_ = 1.65, *P* = 0.22 for the interaction between Q/R editing and the mutation. **e**) Pooled data (as in c) for the fractional steady state current (*I*_SS_). Two-way ANOVA indicated: *F* _1, 17_ = 65.37, *P* < 0.0001 for Q/R editing; *F* _1, 17_ = 28.37, *P* < 0.0001 for the mutation; *F* _1, 17_ = 14.93, *P* = 0.0012 for the interaction between Q/R editing and the mutation. Indicated *P* values are from pairwise Welch *t* -tests.

### A kinetic model incorporating conducting desensitized receptors

We next considered whether the presence of conducting desensitized receptors could account quantitatively for our observations. To examine this, we incorporated such states into a modified version of a kinetic scheme we used previously to describe AMPAR/TARP concentration-dependent behaviors^25^ (Scheme 1; **Fig. 5a)**, and attempted to mimic the macroscopic kinetics and NSFA of GluA2(R)/γ-2 by varying the rate constants and conductances. From six patches in which activation, deactivation and desensitization were all examined, we generated global average waveforms and current-variance plots for each condition (**Fig. 5b-d**). The inclusion in our scheme of conducting desensitized states allowed simultaneous modeling of all kinetic and current-variance data.

**Figure 5.**
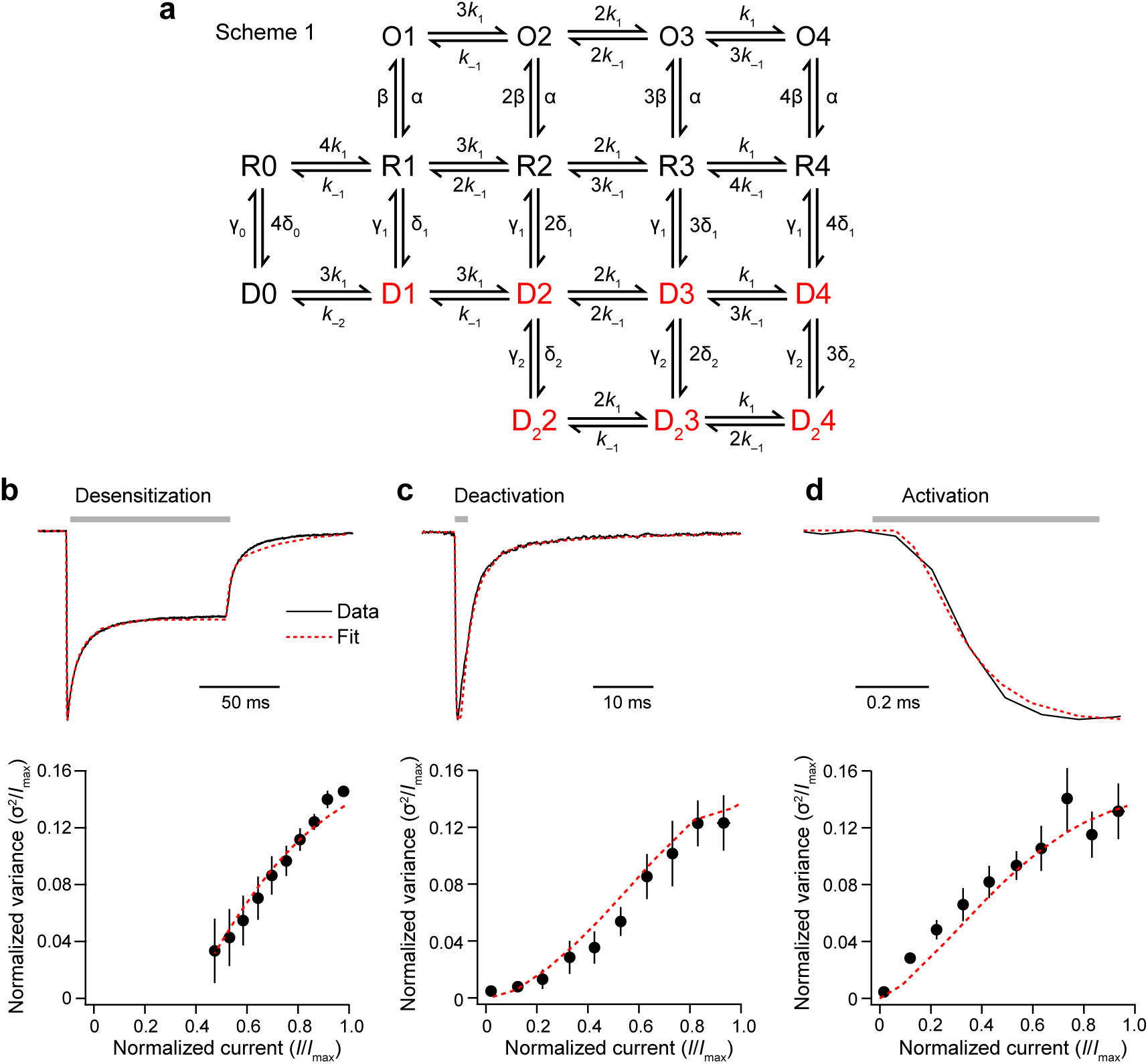
A kinetic scheme including conducting desensitized states can mimic GluA2(R)/γ-2 behavior. **a**) Scheme 1 is a modified form of a previously proposed kinetic model^25,44^. States which can conduct are red. Open states (O1-O4) and occupied desensitized states (D1-D4, D_2_2-D_2_4) both have independent conductances that are occupancy-dependent. **b-d**) Global averaged GluA2(R)/γ-2 records and variance data (below) for desensitization, deactivation and activation (10 mM glutamate – gray bars). Using a single set of rate constants and conductances, the model closely mimics all six measures (dashed red lines): *k* _1_ = 1.3 * 106 M^−1^ s^−1^, *k* _−1_= 350 s^−1^, α = 3,100 s^−1^, β = 1,000 s^−1^, γ_1_ = 88 s^−1^, d_1_ = 110 s^−1^, γ_2_ = 36 s^−1^, d_2_ = 39 s^−1^, γ_0_ = 8 s^−1^, d_0_ = 0.48 s^−1^, *k* _−2_ = 870 s^−1^, conductance of fully-occupied open state (O4) = 3.9 pS, conductance of fully-occupied desensitized state (D4 and D_2_4) = 670 fS.

The model yielded estimated conductances of 3.9 pS for the fully open state and 670 fS for the fully occupied desensitized state of GluA2(R)/γ-2. As the steady-state occupancy of desensitized receptors (D1– D4, D_2_2–D_2_4 combined: 90%) was much higher than that of the open receptors (O1–O4 combined: 6%), the latter contributed just 27% of the steady-state current. The low conductance and high steady-state occupancy of desensitized channels predicted by the model (**Fig. 5b**) can fully explain the unusually low variance of the large steady-state current and, there-fore, the right-shifted current-variance relationship produced from the macroscopic desensitizing current (**Fig. 1**). The model indicated that the proportion of current carried by desensitized and non-saturated receptors increased during deactivation (from 11% and 6%, respectively, at the peak, to 25% and 77% at mid-decay). The combination of these factors explains the rapid fall-off in deactivation variance (**Supplementary Fig. 2** and **Fig. 5c**). Thus, the existence of conducting desensitized states is able to account for many of our observations with homomeric GluA2(R) complexes.

### Functional cross-linking of the conducting desensitized state

Recent work on the structural basis of desensitization has shown that the variety of conformations adopted by the desensitized LBD layer is greatly diminished when the full-length AMPAR is co-assembled with auxiliary subunits^16,19,28,29^. In the presence of γ-2 the LBD dimers of desensitized GluA2 favor a ‘relaxed dimer’ conformation, with the upper D1-D1 interfaces ruptured and the lower D2 domains more closely apposed, allowing channel closure^16^. We reasoned that conducting desensitized receptors could enable us to determine whether this conformation (previously revealed through crystallography and cryo-EM) can indeed be adopted by AMPARs in the plasma membrane. Specifically, we predicted that if the steady-state current of GluA2(R) did indeed reflect ion flow through desensitized receptors in the relaxed dimer conformation, then if trapped in this conformation the receptors should maintain their conductance. To test this, we introduced cysteines at sites on the central axis of the D2-D2 dimer interface, cross-linking of which has previously been shown to inhibit channel opening by trapping the receptor in desensitized-like states. Thus we compared S729C, cross-linking of which permits the relaxed dimer conformation^16,18,45^, with G724C^46^ which we predicted would not accommodate the relaxed dimer conformation when crosslinked (**Supplementary Fig. 4**). In each case, we examined the effect of cross-linking on the steady-state GluA2(R)/γ-2 current.

We first simulated currents from wild-type GluA2(R)/γ-2 receptors (Scheme 1), and compared these with simulated currents expected from receptors occupying only desensitized states (Scheme 2; **Fig. 6a,b**). These simulations predicted that if cross-linking trapped receptors in the native conducting desensitized state, it would change the glutamate response to a purely non-decaying steady-state current of reduced size.

**Figure 6.**
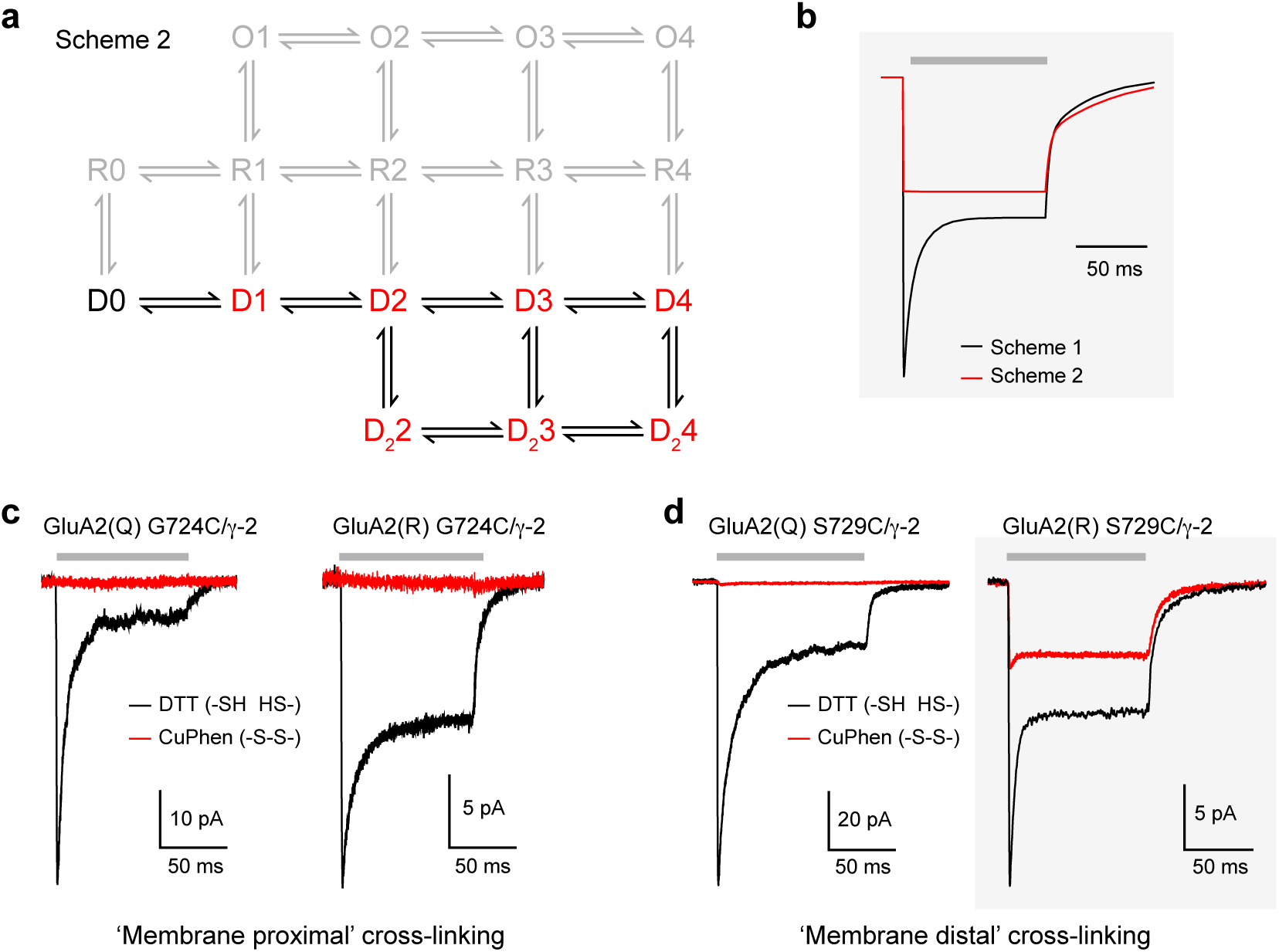
A model with access to only desensitized states predicts behavior of cross-linked GluA2(R) S729C/γ-2. **a**) Scheme 2 is modified from Scheme 1 (Fig. 5a) and assumes that, following S729C cross-linking, the receptor can occupy only desensitized states (excluded states are shown in gray). **b**) Simulated responses to 10 mM glutamate (gray bar) using Scheme 1 to mimic the non-cross-linked condition and Scheme 2 to mimic the effect of cross-linking. **c**) Representative currents at –60 mV activated by 10 mM glutamate (gray bar) from GluA2(Q) G724C/γ-2 and GluA2(R) G724C/γ-2 in DTT (black) or CuPhen (red). Note that, for both forms, currents are fully inhibited following cross-linking by 10 μM CuPhen. **d**) Representative responses from individual patches demonstrate that following cross-linking by 10 μM CuPhen, GluA2(Q) S729C/γ-2 currents are inhibited, while GluA2(R) S729C/γ-2 currents show minimal desensitization and continue to display a large steady-state current, as predicted in a. Gray boxes (b and d) highlight the similarity of modeled currents and recorded GluA2(R) S729C/γ-2 currents.

To investigate the functional effects of cysteine cross-linking, and test these predictions, we examined the sensitivity of both unedited and edited GluA2 G724C/γ-2 and S729C/γ-2 receptors to the oxidizing agent CuPhen^45^. Application of CuPhen to both editing forms of GluA2 G724C/γ-2 receptors caused rapid inhibition of glutamate-evoked currents, resembling the reported effects of cross-linking on TARP-free receptors^46^ (**Fig. 6c**; **Supplementary Fig. 5a**). The peak and steady-state currents generated by GluA2(Q) S729C/γ-2 were also completely inhibited by CuPhen (**Fig. 6d**; **Supplementary Fig. 5b**). By contrast, cross-linking of edited GluA2(R) S729C/γ- 2 abolished the ‘peak’ current, but not the steady-state current (**Fig. 6d**; **Supplementary Fig. 5b**)–a result consistent with the predictions of Scheme 2. Similar results were also seen when GluA2(R) S729C was expressed with γ-8 or GSG1L (**Supplementary Fig. 5c**). Overall, these results demonstrate that ion permeation through desensitized channels allows possible LBD conformations of the desensitized wild-type receptor to be probed using functional cross-linking. Of the two LBD conformations we examined, only S729C, which can accommodate the ‘relaxed dimer’ state when cross-linked, behaved in a manner resembling that of the wild-type desensitized channel.

### Antagonist-bound cross-linked GluA2 LBD structures

Our functional data demonstrate that cross-linked GluA2(R) S729C/γ-2 receptors retain sensitivity to glutamate, implying that upon agonist binding they can undergo structural change which affects the channel gate. To understand the molecular basis of this, we determined the crystal structure of cross-linked S729C ligand binding cores in apo-like conformations bound to the competitive antagonists NBQX (diffracted to 1.8 Å resolution) or ZK200775 (diffracted to 2.0 Å) (**Fig. 7a**; **Supplementary Fig. 6**). AMPAR gating is driven by the separation of the D2-M3 linker regions following agonist binding^12,16,17^. Thus, binding of glutamate is known to increase the distance between the α-carbons of Proline 632 pairs at the base of the D2 lobe of the GluA2 LBD (**Fig. 7a**)^12,15,47^. For the cross-linked S729C mutant LBD bound with NBQX (S729C_NBQX_) the Proline 632 separation was 22.8 Å (similar to that seen with S729C_ZK_, 22.6 Å). This is less than the separation we calculate for the glutamate-bound mutant LBD (S729C_glu_, 26.4 Å)^18^. The relative separation of Pro632 residues suggests that despite being constrained by the cross-link, glutamate binding to GluA2(R) S729C can trigger relative movements of the lower LBDs which may induce tension in the M3-D2 linkers sufficient to allow ion flow (**Fig. 7b**).

**Figure 7.**
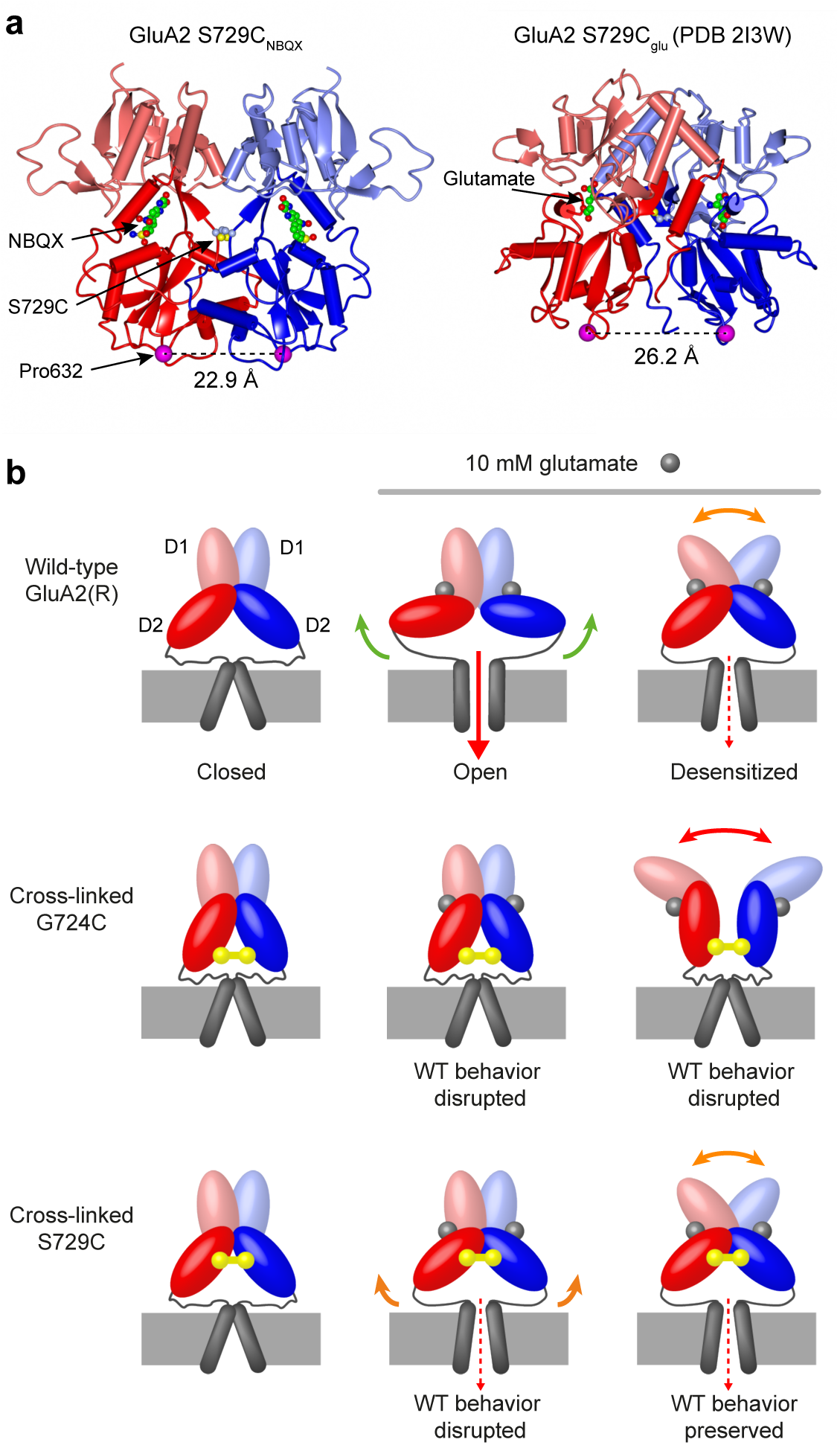
Ligand-dependent differences in cross-linked S729C LBD structures suggest a model of gating for desensitized GluA2(R). **a**) Left, crystal structure of the dimeric GluA2 S729C ligand binding core in the presence of NBQX, with the upper (D1, pale) and lower (D2, dark) lobes of each monomer (red and blue) distinguished by shading. The structure is viewed perpendicular to the axis between the Cα atoms of Pro632 (magenta spheres). Right, crystal structure of the GluA2 S729C ligand binding core in the presence of glutamate (S729C_glu_; PDB: 2I3W)^18^. The Pro632 separation seen in the presence of glutamate (right) is > 3Å greater than that seen with NBQX (left), and we suggest that this is sufficient to promote channel opening despite S729C cross-linking. **b**) Cartoon representing possible conformations of the GluA2(R) LBD dimer and pore in our functional cross-linking recordings. Non-cross-linked GluA2(R) channels bind glutamate (gray sphere), closing the clamshell LBDs and opening the pore to the full open channel conductance. Desensitization does not fully close the pore. As determined in the presence of γ-2, cross-linking of the G724C mutant (yellow) does not allow the action of agonist binding to be communicated to the pore in any state, and disrupts the normal dimeric conformation of desensitized receptors. Cross-linking of the S729C mutant (yellow) is not compatible with the full open state, but the channel can adopt the normal desensitized conformation, meaning that (as with the non-cross-linked receptor) the pore is not closed.

We also crystallized isolated cross-linked G724C LBDs in both the apo-like ZK200775-bound form (G724C_ZK_) and desensitized-like glutamate-bound forms in two different space groups (G724C_glu_ Forms A and B; **Supplementary Fig. 7**). As we anticipated, G724C_ZK_ demonstrated reduced Pro632 separation (19.2 Å) compared to the antagonist-bound S729C forms, in keeping with the greater constraints on LBD separation imposed by G724C. However, unlike glutamate-bound S729C^18,45^, neither form of G724C_glu_ adopted a relaxed dimer conformation (**Fig. 7b**). Instead they displayed a non-biological conformation, with a large rotation and total loss of the dimer interface which can not be accommodated within the intact receptor. Thus, the Pro632 separation in the intact, cross-linked GluA2 G724C receptor could not be meaningfully assessed.

## Discussion

Our experimental findings have important implications for the understanding of AMPAR desensitization. They challenge the dogma that the desensitized AMPAR pore is, by definition, closed, while providing direct functional evidence concerning the conformation of the ligand binding domains of desensitized receptors. First, our NSFA, single-channel measurements and kinetic modeling all support the unexpected conclusion that atypical currents generated by GluA2(R)/γ-2 are mediated by ‘conducting desensitized’ receptors, which retain around one sixth of the maximal conductance of open channels. While a conducting desensitized state has previously been described for a mutant, homomeric, α7 nicotinic acetylcholine receptor^48^, to our knowledge this is the first report of such behavior for a ligand-gated ion channel formed from wild-type subunits. Second, the presence of a conducting desensitized state of GluA2(R) provided us with a novel functional read-out to probe the likely arrangement of LBDs within desensitized AMPARs. By comparing currents from GluA2(R)/γ-2, GluA2(R)/γ-8 and GluA2(R)/GSG1L receptors with those from constrained cross-linked cysteine mutants^18,46^, we have established that the ‘relaxed’ dimer conformation identified in structural studies^18^, is indeed representative of desensitized AMPARs within the plasma membrane.

What is the evidence that the steady state current does indeed arise from desensitized AMPARs? The steady-state glutamate-evoked currents we recorded with GluA2(R)/γ-2 were much larger than those of other AMPAR/auxiliary subunit combination we have examined previously^34–38^. While one might reasonably suppose that such a large steady-state current could arise simply from a reduced level of receptor desensitization, a number of the GluA2(R)/γ-2 properties we have identified suggest this is not the case. Importantly, the rates of entry into, and recovery from, desensitization for GluA2(R)/γ-2 were similar to those for GluA2(Q)/γ-2, which had a much smaller steady-state current. Moreover, a reduced extent of desensitization would not give rise to the rightward shift in the current-variance relationships that we observed. By contrast, the presence of a substantial steady-state current of unusually low variance would produce such a shift. Thus, the steady-state current could be generated by low-conductance channels with a high open probability. Furthermore, resolvable (picosiemens) openings were common at the onset of the current, but occurred only rarely during steady-state, supporting the view that these contribute little to the large steady-state component. Finally, AMPAR mutations (S754D and S729C) that enhanced desensitization essentially eliminated steady-state currents of GluA2(Q)/γ-2 receptors, but had much less effect on steady-state currents of GluA2(R)/γ-2 receptors. Consequently, our data strongly suggest that a large fraction of the GluA2(R)/γ-2 steady-state current arises from conducting desensitized channels.

How do our functional data fit with recently published AMPAR structures? Cryo-EM analysis has indicated that the pore diameter at the M3 gate of agonist-bound desensitized GluA2(R)/γ-2 is less than that of the activated receptor^16^ and similar to that of the closed (antagonist-bound) receptor^33^. Thus, our demonstration of conducting desensitized states could be seen to present something of a paradox. The single closed structure for desensitized GluA2(R)/γ-2 (Ref. 16) indicates that ‘open’ desensitized states were not present under the conditions used for cryo-EM analysis or, if present, were either too heterogeneous to allow reconstruction, or perhaps too similar in structure to ‘closed’ de-sensitized states to be classified distinctly. At the same time, although our kinetic and current-variance data could be adequately mimicked by Scheme 1 without the inclusion of closed desensitized receptors, our data do not preclude their existence. For simplicity, we assigned a conductance to all occupied desensitized states, but models including both closed and conducting desensitized states could also broadly reproduce our functional data. Nonetheless, the magnitude of the steady-state current (relative to the peak current) limits how many closed desensitized channels are likely to be present in our recordings. Of note, rapid transitions between these states would be required to account for the absence of clear channel closures from steady-state in our single-channel records. Taken together, it is certainly possible that both closed and open desensitized re-ceptors are present in cryo-EM conditions, in which closed desensitized channels might predominate, as well as in our recordings, in which conducting desensitized channels are clearly prevalent.

In our model of GluA2(R)/γ-2 gating, assigning fully-occupied desensitized channels a conductance of 670 fS provided a good approximation to both our kinetic and noise data. While currents mediated by GluA2(R), GluA2(R)/γ-8 and GluA2(R)/GSG1L also bore all the hallmarks of conducting desensitized channels, for these combinations the deviations from conventional parabolic current-variance relationships were less dramatic, and the editing-dependent increases of steady-state currents were smaller. Furthermore, when co-expressed with γ-8 or GSG1L, the cross-linked GluA2(R) S729C mutant displayed a smaller residual steady-state current than that seen when it was co-expressed with γ-2. One possible explanation for these differences is that for GluA2(R) receptors the rank-order of desensitized channel conductance is γ-2 > γ-8 > no auxiliary > GSG1L. An alternative possibility is that this reflects the presence of both conducting and non-conducting desensitized states. In this latter scenario, our macroscopic data could be explained by γ-2-associated receptors spending a greater proportion of time than GluA2(R), GluA2(R)/γ-8 or GluA2(R)/GSG1L in conducting desensitized states, relative to closed desensitized states.

Recently, a chimeric AMPAR/KAR construct (ATD and LBD of GluK2 with TM and C-tail of GluA2) has been shown to exhibit a large ‘leak’ current when co-expressed with TARPs^49^. Remarkably, when these chimera/TARP combinations were exposed to glutamate the currents decreased, suggesting that, despite conducting in the absence of agonist, desensitization could still cause closure of the channel^49^. The existence of a leak current in the presence of a TARP was taken to indicate that TARPs disrupt the ligand-free ‘closed’ state of the receptor, leading to spontaneous channel opening. Although we found no evidence that edited GluA2 receptors open spontaneously when expressed with a TARP, the fact that they conduct when desensitized – and the fact that the magnitude of the steady-state current is greatest when γ-2 is present – supports the view that TARPs can also disrupt channel closure in the desensitized state, in a manner that is Q/R-editing dependent.

How might Q/R editing render the desensitized GluA2 receptor ion permeable? The switch from a neutral glutamine to a positively changed arginine alters both the charge and volume of the sidechain at the Q/R site. While the orientation of the arginine side chains in the desensitized state has yet to be resolved, in the activated structure of the GluA2(R)/γ-2 receptor^16^ (and in closed heteromeric GluA1/2(R)_γ-8; Ref 27) they project away from the cytoplasm and towards the gate. Structural modelling of the equivalent arginines in the highly homologous homomeric edited kainate receptor GluK2(R) suggests a similar outward projection of the side chains, resulting in an interaction between the pore loop and the M3 helix proximal to the gate that modifies the channel pharmacology^50^. If such interactions are present in GluA2(R), these might modify the rearrangement of the channel gate following desensitization, thereby hampering full channel closure. Alternatively, charge-charge repulsion between arginines at the level of the selectivity filter could compromise the narrowing of the pore and may be sufficient, on its own, to account for the ion flow through desensitized channels.

There is general agreement that desensitization-induced AMPAR pore closure is caused by LBD rearrangements which allow the base of the D2 lobes to assume positions similar to those of the apo/inactivated form, thereby releasing the tension exerted by the M3-S2 linkers on the M3 helix caused by glutamate binding^16,19^. In the absence of auxiliary proteins there is considerable heterogeneity in the LBD and ATD layers of desensitized AMPARs^28,29^. However, when associated with γ-2 or GSG1L, AMPAR LBD dimers show increased stability of a single ‘relaxed dimer’ conformation^16,18,19^. The fact that GluA2(R) S729C, co-expressed with auxiliary subunits and trapped in the relaxed dimer conformation by cross-linking, exhibited properties consistent with those of wild-type channels, demonstrates that this conformation does indeed mimic the behavior of the native desensitized receptor in the plasma membrane. By contrast, functional cross-linking of a different mutant GluA2(R) G724C/γ-2 – which did not assume a relaxed LBD dimer when crystallized – trapped these receptors in a non-conducting state, indicating that this conformation must be distinct from that of native desensitized AMPARs. It is noteworthy that despite the constraint on LBD movement imposed by cross-linking at S729C, the current produced by cross-linked GluA2(R)/γ-2 was glutamate-dependent, suggesting agonist binding produces changes within the constrained LBD layer sufficient to influence the pore. Consistent with this, dimeric Pro632 separation within S729C ligand binding core crystals is increased in the presence of glutamate^18^, compared to the separation we observed in the presence of competitive antagonists. While this suggests that agonist binding can potentially exert a small degree of tension on the LBD-TM linkers within the cross-linked receptors, the Pro632 measure provides only a one dimensional approximation of a complex three dimensional process. Future cryo-EM analysis of full-length GluA2 S729C may provide valuable further information on the complex dynamics of desensitized receptors.

Does the conducting desensitized state of homomeric GluA2(R) contribute to neuronal or glial signaling? Neurons and glia normally express multiple AMPAR subunit isoforms. When GluA2 is coexpressed with other subunits (in the absence of auxiliary proteins) the formation of GluA2 homomers is strongly discriminated against, in favor of GluA2-containing heteromers^51,52^. Nonetheless, trafficking of homomeric GluA2(R) is enhanced if the receptors are unedited at the secondary (R/G) editing site^53^, and we (and others^54,55^) have demonstrated that the presence of γ-2 allows robust heterologous expression of functional GluA2(R) homomers. Of note, glutamate-gated channels with femtosiemens conductance have been detected in cerebellar granule cells^56^. Moreover, an immunoprecipitation study that suggested hippocampal AMPARs were predominantly GluA1/2 or GluA2/3 heteromers did not expressly rule out the presence of GluA2 homomers^57^. Additionally, functional GluA2(R) homomers can be trafficked to hippocampal synapses by endogenous TARPs following the conditional deletion of the alleles for GluA1 and GluA3^58^. Thus, while the extent to which TARPs influence the preference of GluA2(R) to heteromerize remains to be determined, it is clear that homomeric GluA2(R) receptors can, in principle, be present at synapses and would conduct even when desensitized.

## Author Contributions

I.D.C., D.S., T.P.M., M.G.G., M.F., and S.G.C-C. designed the experiments. I.D.C., D.S and T.P.M. performed the experiments. I.D.C., D.S., T.P.M., M.G.G. and M.F. analyzed the data. I.D.C., M.G.G., M.F. and S.G.C-C. interpreted the results. I.D.C. and M.F. prepared the figures. I.D.C., M.F. and S.G.C-C. wrote the manuscript with input from all authors.

### Acknowledgements

This work was supported by the MRC (MR/J002976/1 to S.G.C-C. and M.F. and MR/J012998/1 to M.F. and S.G.C-C.), the Wellcome Trust (086185/Z/08/Z to S.G.C-C. and M.F.) and the BBSRC (BB/N015274/1 to M.G.G.). M.G.G. is a Wellcome Trust and Royal Society Sir Henry Dale fellow (104194/Z/14/Z). We thank Ingo Greger for providing the GluA2 LBD expression vector, and Trevor Smart, Duncan Laverty and Ambrose Cole for assistance with crystallography. We thank Mark Mayer for discussion and for comments on an earlier version of the manuscript.

## Data availability

The coordinate and structure factor data for the GluA2 LBD crystals have been deposited in the Protein Data Bank (PDB) with the following accession codes: S729C_NBQX_, 6FQH; S729C_ZK_, 6FQK; G724C_ZK_, 6FQJ; G724C_glu_ Form A, 6FQG; G724C_glu_ Form B, 6FQI.

## Methods

### Heterologous expression

We expressed recombinant AMPAR subunits and TARPs (plus EGFP) in HEK293 cells maintained under standard protocols, as described previously^25^. AMPAR subunit cDNAs (rat) were ‘flip’ splice variants and the GluA2 forms were additionally R/G edited. Point mutations of the GluA2 subunit were produced using standard pcr protocols. AMPAR/TARP combinations were transfected at a cDNA ratio of 1:2. The GluA2_γ-2 tandem consisted of full-length GluA2 and a 9 amino-acid linker (GGGGGEFAT) before the start codon of full-length γ-2. Transient transfection was performed using Lipofectamine 2000 (Life Technologies). Cells were split 12–30 h after transfection and plated on glass coverslips in the presence of 50 μM NBQX (2,3-dioxo-6-nitro-1,2,3,4-tetrahydrobenzo[*f*]quinoxaline-7-sulfonamide; Tocris Bioscience) to avoid AMPAR-mediated toxicity. Electrophysiological recordings were performed 18–48 h later.

### Electrophysiology

Patch-clamp electrodes were pulled from borosilicate glass (1.5 mm o.d., 0.86 mm i.d.; Harvard Apparatus) and fire polished to a final resistance of 8–12 MΩ. For outside-out patches the ‘external’ solution contained 145 mM NaCl, 2.5 mM KCl, 1 mM CaCl_2_, 1 mM MgCl_2_ and 10 mM HEPES, pH 7.3. The ‘internal’ solution contained 145 mM CsCl, 2.5 mM NaCl, 1 mM Cs-EGTA, 4 mM MgATP and 10 mM HEPES (pH 7.3 with CsOH). supplemented with 100 μM spermine tetrahydrochloride (Tocris Bioscience). Currents with a risetime > 500 μs were rejected. For chloride permeability experiments, two CsCl based solutions were used – one with ‘high’ CsCl (145 mM CsCl, 10 mM HEPES, 1 mM CaCl_2_; pH 7.3 with CsOH) and one with ‘low’ CsCl (CsCl reduced to 35 mM and osmolarity adjusted with glucose). For recordings involving cysteine cross-linking, control and agonist solutions were supplemented with 1 mM DTT to reduce disulphide bonds or 10 μM CuCl and 30 μM 1-10-phenanthroline (CuPhen) to promote disulphide formation^45^. The effects of CuPhen were fully reversible by 1 mM DTT. Recordings were made from outside-out patches at 22–25 ^o^C using an Axopatch 200A amplifier (Molecular Devices). Currents were recorded at –60 mV, low-pass filtered at 10 kHz and digitized at 20 kHz, except for recordings to assess activation noise which were digitized at 100 kHz (National Instruments NI USB-6341 interface with Strathclyde Electrophysiology Software WinWCP or Molecular Devices Digidata 1440A interface with pClamp 10 software). Patches with small responses were filtered at 2 kHz to more readily identify single channel openings, and digitized at 10 kHz.

### Rapid agonist application to excised patches

Rapid agonist application was achieved by switching between continuously flowing solutions. Solution exchange was achieved by moving an application tool made from theta glass (Hilgenberg; 2 mm outer diameter, pulled to a tip opening of ∼200 μm) mounted on a piezoelectric translator (Physik Instrumente). At the end of each experiment, the adequacy of the solution exchange was tested by destroying the patch and measuring the liquid-junction current at the open pipette (10–90% rise time typically 150–250 μs).

### Data analysis

Entry into desensitization (100 ms application of 10 mM glutamate) and current deactivation (1 ms) were fitted with the sum of two exponentials using IGOR Pro 6.35 (Wavemetrics) with NeuroMatic^59^. Recovery from steady-state desensitization was measured following a 100 ms equilibrating application of 10 mM glutamate. The recovery of glutamate-activated peak currents was measured following 2–200 ms intervals in control solution.

Records used for single-channel analysis were filtered at 0.5 kHz and individual channel events were selected by eye. Channel openings were analyzed using QuB (ver. 2.0.0.7; http://www.qub.buffalo.edu). The amplitude of the resolved openings was measured from the closing transition (final current level to steady-state current). Measured openings (at –60 mV) were binned by conductance and fitted using a multi-peak Gaussian function (IGOR Pro).

Non-stationary fluctuation analysis (NSFA) was performed on the decaying phase of currents evoked by 1 ms or 100 ms applications of 10 mM glutamate (35–300 successive applications), as previously described^36^. The variance for each successive pair of current responses was calculated and the single-channel current (*i*) and total number of channels were then determined by plotting the ensemble variance (*σ*^2^) against mean current (Ī) and fitting with a parabolic function:

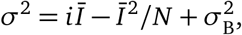

where 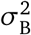 is the background variance^60^. For NSFA of current activation, records were digitized at a high sampling rate (100 kHz) to ensure sufficient numbers of data points from the average record could be grouped into each of the ten amplitude bins. As alignment of traces on their rising phases (as used for deactivation and desensitization records) led to a distortion of activation noise, analysis was instead performed on unaligned traces (from sections of recording in which the time of the current onset was stable; Spearman Stability Analysis, NeuroMatic).

Chloride permeability calculations were made using the equation:

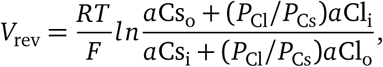

where *V*_rev_ is the reversal potential measured at the peak or steady-state, *P*_Cl_/*P*_Cs_ is the permeability ratio of chloride relative to cesium and *a*Cs and *a*Cl are the activities of the ions in the intracellular (i) and extracellular (o) solutions^43^. *F, R* and *T* have their usual meanings. *a*Cs in the high CsCl solution was extrapolated from tabulated values to be 0.714 (http://www.kayelaby.npl.co.uk/chemistry/3_9/3_9_6.html). *a*Cs in the low CsCl solution was estimated to be 0.824. This value has a small degree of uncertainty, as the effect of glucose – demonstrated to modestly affect *a*Na in NaCl solutions^61^ – is unknown. Our chosen value assumes a similar effect of glucose on the activities of both NaCl and CsCl. To negate the influence of junction potentials, experimentally determined shifts in reversal potentials were compared to the calculated shifts following exchange from the ‘high’ to ‘low’ CsCl solutions (for purely Cs^+^-permeable channels and purely Cl^−^-permeable channels).

### Kinetic Modeling

Kinetic simulations and fits were performed in Scilab 5.5.0. (Scilab Enterprises; http://www.scilab.org) using the Q-matrix method^62^. Rate constant *k*_−2_ was constrained by microscopic reversibility. For each set of rate constants, currents were calculated from the occupancies of all conducting states at given time points multiplied by their unitary current. Noise was calculated using the following equation^63^:

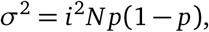

where *N* is the number of channels of unitary current *i* of open probability *p*. The ensemble variance for a channel with multiple subconductances was calculated as the sum of the variances for each state:

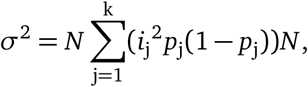

where k is the number of conducting states, j refers to each conducting state, and *i*_j_ and *p*_j_ are its unitary current and occupancy respectively. *N* was re-derived from the experimentally measured peak current, as well as the peak open probability and conductance of each state for each iteration of the fit. Kinetics and noise of desensitization, deactivation and activation, across 6 patches from which all properties could be measured, were normalized, averaged and fit using Scheme 1 (**Fig. 5**). Kinetic data were parsed (to 35 data points) to make computation manageable.

### Expression and purification of ligand binding cores

Cysteine mutants of the GluA2 flop S1S2 binding core with an N-terminal octahistidine tag in the pET22b vector^12^ were produced using standard PCR protocols. Following transformation of Origami B cells, high levels of protein were expressed by induction with 0.5 mM IPTG when the cells had reached OD_600_∼1.2. The cells were harvested by centrifugation following overnight incubation at 20°C. Harvested cells were washed once with PBS and resuspended in His-Trap binding buffer (50 mM Tris, pH 8; 150 mM NaCl; 20 mM imidazole; protease inhibitor cocktail (Roche) and 50 μM NBQX to displace glutamate and promote dimerization). Resuspended cells were treated with lysozyme for 30 mins at 4°C and cell membranes were disrupted by sonication and removed by centrifugation. Samples were filtered (0.45 μm) to remove residual cellular debris and loaded onto a His-Trap Column (GE Healthcare) at 4°C. Protein was eluted using HisTrap elution buffer (50 mM Tris pH 8; 150 mM NaCl; 300 mM imidazole) and aliquots containing the highest concentrations of dimers were pooled for further purification. Parallel reducing and non-reducing SDS-PAGE and Coomassie blue staining established that ligand binding cores preferentially formed cross-linked dimers with no need for exogenous oxidization. Protein was concentrated using 10 kDa concentrators before exchanging into column buffer (50 mM Tris, pH 8; 150 mM NaCl). Histidine tags were cleaved using thrombin. A final purification step (in column buffer) was performed with a size exclusion column (Superdex 200; GE Healthcare, Little Chalfont, UK). Final purified AMPAR LBDs were concentrated to 2–7 mg/ml.

### Protein crystallography

Crystallization was achieved using sitting drop vapor diffusion at 16°C. All crystals appeared within 72 hours, and were harvested after 1–2 weeks. For each crystal, precipitant solutions, and additives for cryoprotection prior to freezing, were as follows:

- GluA2 S729C_NBQX_: 0.1 M tri-sodium citrate pH 5.5, 20% PEG 3000. Supplemented with 15% glycerol for cryoprotection.
- GluA2 S729C_ZK_/GluA2 G724C_ZK_: 0.2 M ammonium chloride, 20% PEG 3350, 10 μM ZK200775. Supplemented with 15% glycerol for cryoprotection.
- GluA2 G724C_glu_ Form A: 1 M lithium chloride, 0.1 M citric acid pH 4.0, 20% (w/v) PEG 6000, 30 mM glutamate. Supplemented with 20% glycerol for cryoprotection.
- GluA2 G724C_glu_ Form B: 0.16 M calcium acetate, 0.08 M sodium cacodylate pH 6.5, 14.4% (w/v) PEG 8000, 20% (v/v) glycerol, 1 mM glutamate. No additives necessary for cryoprotection.

Diffraction data were collected at Diamond Light Source beamlines I04 and I24, and at ESRF ID30B (see **Supplementary Table 1**). Diffraction data were initially processed using xia2^64^ and AIMLESS^65^. Initial molecular replacement was performed using Phaser^66^ and structures were refined using PHENIX^67^ and *Coot*^68^. Structures G724C_ZK_, S729C_NBQX_ and S729C_ZK_ were solved using the ZK200775 bound wild-type LBD (PDB 3KGC)^13^ as the search model. Both forms of G724C_glu_ were solved using the glutamate bound wild-type LBD (PDB 1FTJ)^12^ as the search model. Cysteines were modeled into cryo-EM structures of GluA2(R)/γ-2 in the activated and desensitized forms using PyMOL (The PyMOL Molecular Graphics System, Version 2.0 Schrödinger, LLC.), and the separation of the sulphur atoms was determined. The separation of Cα Pro632 atoms in LBD structures was also measured using PyMOL.

### Data presentation and statistical analysis

Summary data are presented in the text as mean ± s.e.m. (from *n* patches). Comparisons involving two data sets only were performed using a two-tailed paired *t*-test or two-tailed unpaired Welch two-sample *t*-test that does not assume equal variance (normality was not tested statistically, but gauged from quantile-quantile plots and/or density histograms). Comparisons of multiple conditions were performed using two-sided Welch two-sample *t*-tests with Holm’s sequential Bonferroni correction. When comparing Q and R edited forms of AMPARs, analyses were performed using two-way analysis of variance (Welch heteroscedastic *F*-test) followed by pairwise comparisons using two-tailed Welch two-sample *t*-tests. Exact *P* values are presented to two significant figures, except when *P* < 0.0001. Statistical tests were performed using R (version 3.5.2, the R Foundation for Statistical Computing, http://www.r-project.org/) and R Studio (version 1.2.1303, RStudio). No statistical test was used to predetermine sample sizes; these were based on standards of the field. No randomization was used. A full list of statistical analyses is provided in **Supplementary Table 2**.

