## Supplementary File for "Homomeric Q/R edited AMPA receptors conduct when desensitized"

### Supplementary Information

1. **Supplementary Figure 1. Current-variance relationships of desensitization for different AMPAR constructs and voltages.**
2. **Supplementary Figure 2. Current-variance relationships of deactivation for edited AMPARs with $\gamma$-2.**
3. **Supplementary Figure 3. GluA2(R)/$\gamma$-2 receptors at both peak and steady-state mediate negligible $Cl^-$ flux.**
4. **Supplementary Figure 4. Predicted minimum separations of sulfur atoms for mutant cysteines modeled into quisqualate-bound GluA2/$\gamma$-2 structures.**
5. **Supplementary Figure 5. Globally averaged currents showing the effects of cross-linking for GluA2 G724C and S729C mutants.**
6. **Supplementary Figure 6. Electron densities of ligands and disulfide bonds for $S729C_{NBQX}$ and $S729C_{ZK}$.**
7. **Supplementary Figure 7. G724C cross-linking disrupts the relaxed dimer structure of LBD.**
8. **Supplementary Table 1. Data collection and refinement statistics.**
9. **Supplementary Table 2. Details of statistical analyses.**

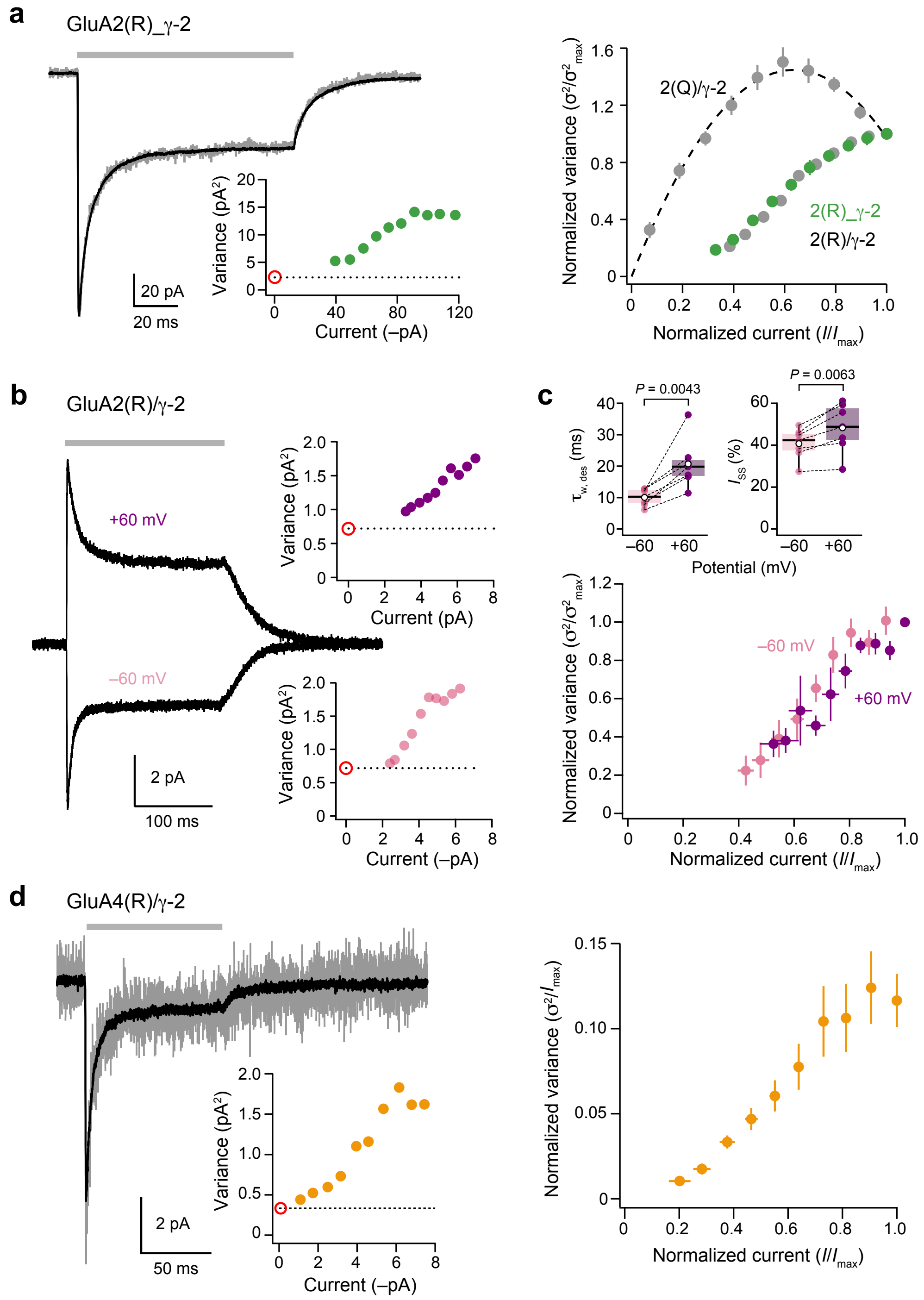

**Supplementary Figure 1. Current-variance relationships of desensitization for different AMPAR constructs and voltages**. **a**) Representative current (gray; average in black) from outside-out patch taken from a tandem GluA2(R)_γ-2-expressing HEK293 cell, evoked by 100 ms application of 10 mM glutamate at –60 mV (gray bar). Inset: current-variance relationship for the same patch (dotted line indicates background variance and open red circle indicates expected origin). Note that the current-variance relationship cannot be fitted with a parabola passing through the origin. Right panel shows doubly normalized and averaged current-variance relationships for desensitization of GluA2(R)_γ-2 (green circles; $n$ = 8). Also shown, for comparison, are corresponding relationships for GluA2(Q)/γ-2 and GluA2(R)/γ-2 (gray circles, from Fig. 1). Error bars denote sems. **b**) Averaged GluA2(R)/γ-2 currents evoked by 200 ms applications of 10 mM glutamate (gray bar) at +60 and –60 mV. Insets: current-variance relationships for the same responses (displayed as in a). Note that in neither case can the current-variance relationship be fitted with a parabola passing through the origin. **c**) Upper panels show pooled data for kinetic properties ($n$ = 7 for both). Box-and-whisker plots as in Fig. 1c, but with dashed lines linking paired measurements. Indicated $P$ values from paired $t$-tests). Lower panel shows doubly normalized and averaged current-variance relationships for desensitization of GluA2(R)/γ-2 at +60 and –60 mV ($n$ = 5 and 6, respectively). Error bars denote sems. **d**) Representative GluA4(R)/γ-2 current (gray, average in black) evoked by 100 ms application of 10 mM glutamate at –60 mV. Inset: current-variance relationship for the same patch (displayed as in a). Note that the current-variance relationship cannot be fitted with a parabolic relationship passing through the origin. Right panel shows normalized and averaged current-variance relationships for desensitization of GluA4(R)/γ-2 ($n$ = 6). Error bars denote sems.

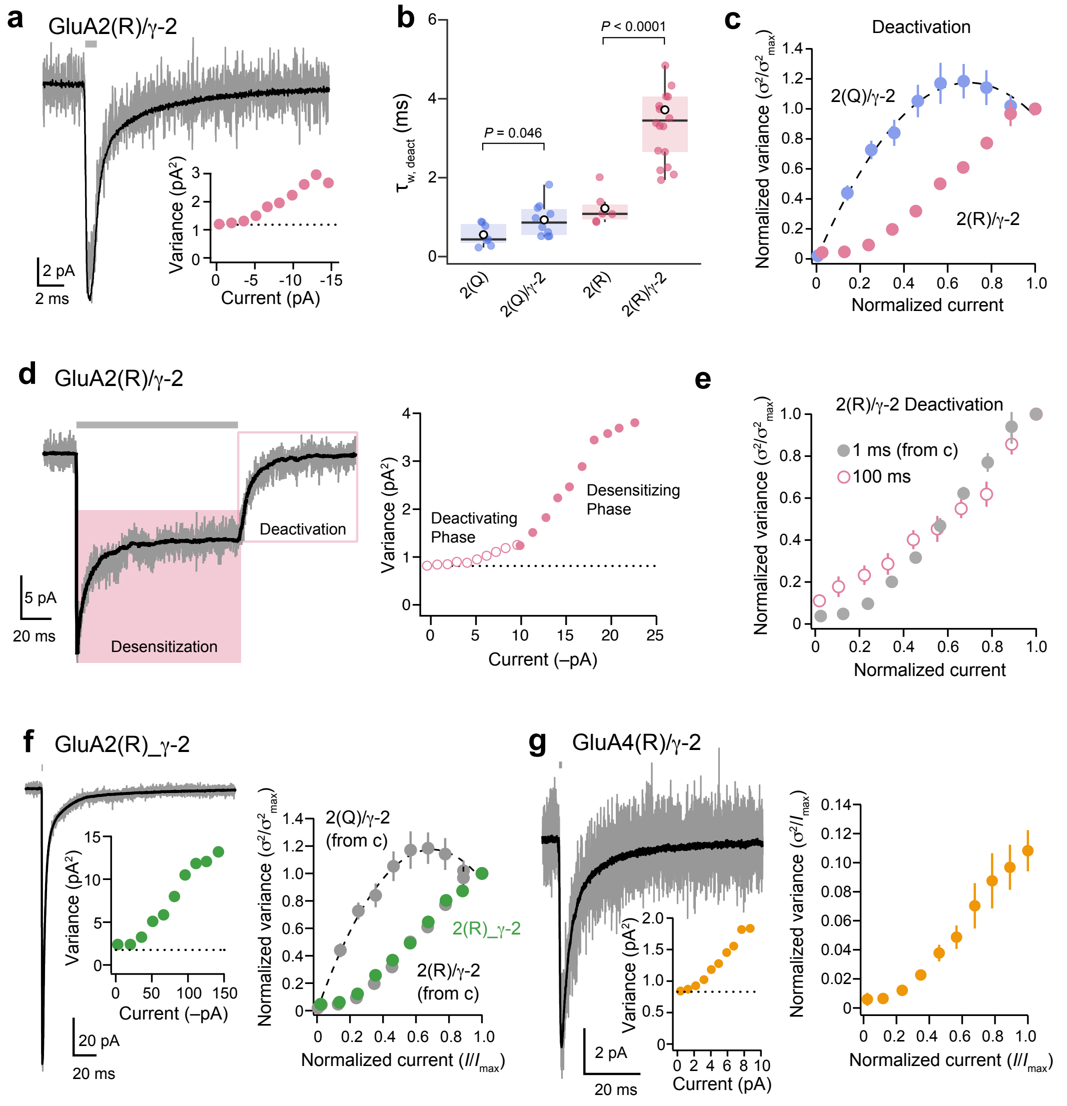

**Supplementary Figure 2. Current-variance relationships of deactivation for edited AMPARs with γ-2.** **a**) Representative GluA2(R)/γ-2 current (gray, average in black) evoked by 1 ms application of 10 mM glutamate at –60 mV (gray bar) to an outside-out patch from a transfected HEK293 cell. Inset: current-variance relationship for the same patch (dotted line indicates background variance). Note that the data can not be fitted with a parabola. **b**) Pooled deactivation kinetics for GluA2(Q) and (R) with and without γ-2 (for Q $n$ = 10 and 7; for R $n$ = 18 and 6, respectively). Box-and-whisker plot as in Fig. 1c. Two-way ANOVA indicated: $F_{1,37}$ = 34.70, $P$ < 0.0001 for Q/R editing; $F_{1,37}$ = 17.47, $P$ = 0.00017 for γ-2; $F_{1,37}$ = 20.64, $P$ < 0.0001 for the interaction between Q/R editing and γ-2. Indicated $P$ values are from pairwise Welch $t$-tests. **c**) Doubly normalized and averaged current variance relationships of deactivation for GluA2(Q)/γ-2 ($n$ = 7) and GluA2(R)/γ-2 ($n$ = 18). Error bars denote sems. **d**) A representative GluA2(R)/γ-2 current evoked by a 100 ms application of 10 mM glutamate at –60 mV. NSFA was performed on both the desensitizing phase (pink shaded region; as performed in Fig. 1), and on the deactivating phase following glutamate removal (open pink box). Right panel shows current-variance relationship for both phases of the current. **e**) Doubly normalized current-variance relationships for deactivation of GluA2(R)/γ-2 (1 ms, duplicated from c; 100 ms, $n$ = 11). Note the similarity in shape of the relationships. Error bars indicate sems. **f**) Deactivation current and current-variance relationship for the tandem construct GluA2(R)_γ-2. Representative trace in gray and average in black. The right hand panel shows pooled doubly normalized current and variance ($n$ = 5) compared with separately expressed subunits (from c). Error bars indicate sems. **g**) As for f, but for GluA4(R)/γ-2 deactivation ($n$ = 5). Error bars indicate sems.

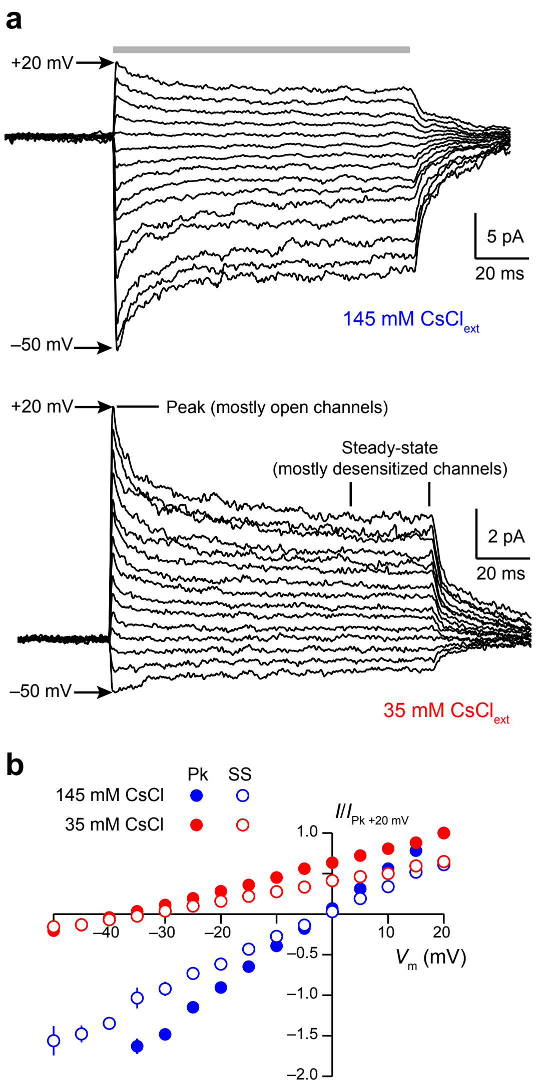

**Supplementary Figure 3. GluA2(R)/γ-2 receptors at both peak and steady-state mediate negligible Cl$^-$ flux**. **a**) Representative GluA2(R)/γ-2 currents evoked by 100 ms applications of 10 mM glutamate (gray bar) to an outside-out patch from a transfected HEK293 cell (at potentials between –50 and +20 mV; Δ5 mV). Currents were recorded in external solution containing either 145 mM CsCl, or 35 mM CsCl. **b**) Mean current-voltage relationships (normalized to the peak current at +20 mV) for the high CsCl solution (blue) and the low CsCl solution (red) ($n$ = 7). Error bars denote sems. Note the similar shift in reversal potential of both the peak (Pk) and steady-state (SS) components.

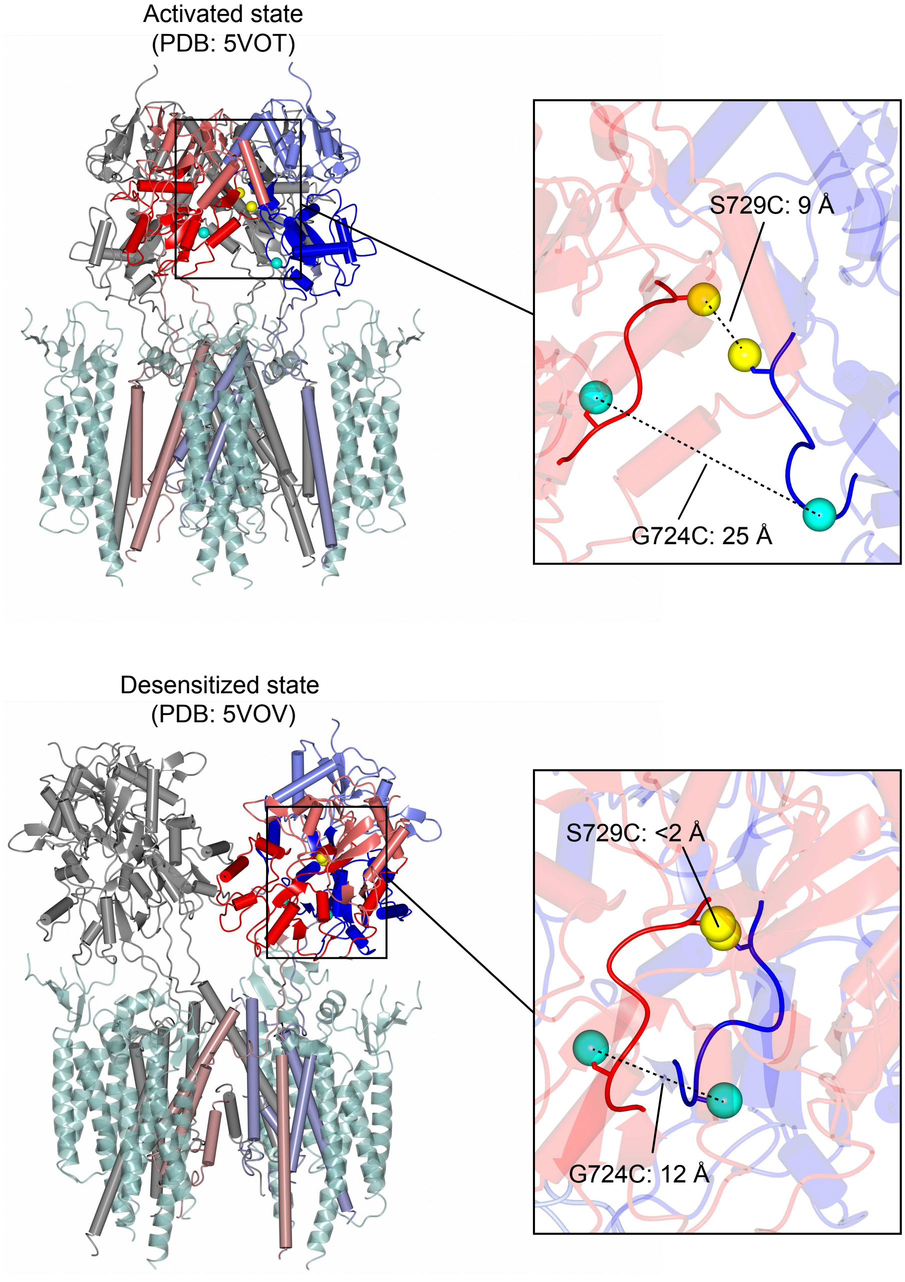

**Supplementary Figure 4. Predicted minimum separations of sulfur atoms for mutant cysteines modeled into quisqualate-bound GluA2/$\gamma$-2 structures**. Cysteine substitutions at Ser729 and Gly724 were modeled into cryo-EM structures of GluA2(R)/$\gamma$-2 in the activated state (upper) and desensitized-like state (lower) (from Ref. 16). Subunits 'A' and 'D' are shown in red and blue, respectively, with subunits 'B' and 'C' (gray) and all four $\gamma$-2 subunits (sea green, transparent). Expanded sections highlight the sulfur atoms of the modeled cysteines at positions 724 (cyan) and 729 (yellow), with the calculated separations. The modeling predicts that a disulfide bond (2 Å) can only be accommodated by desensitized GluA2(R) S729C/$\gamma$-2.

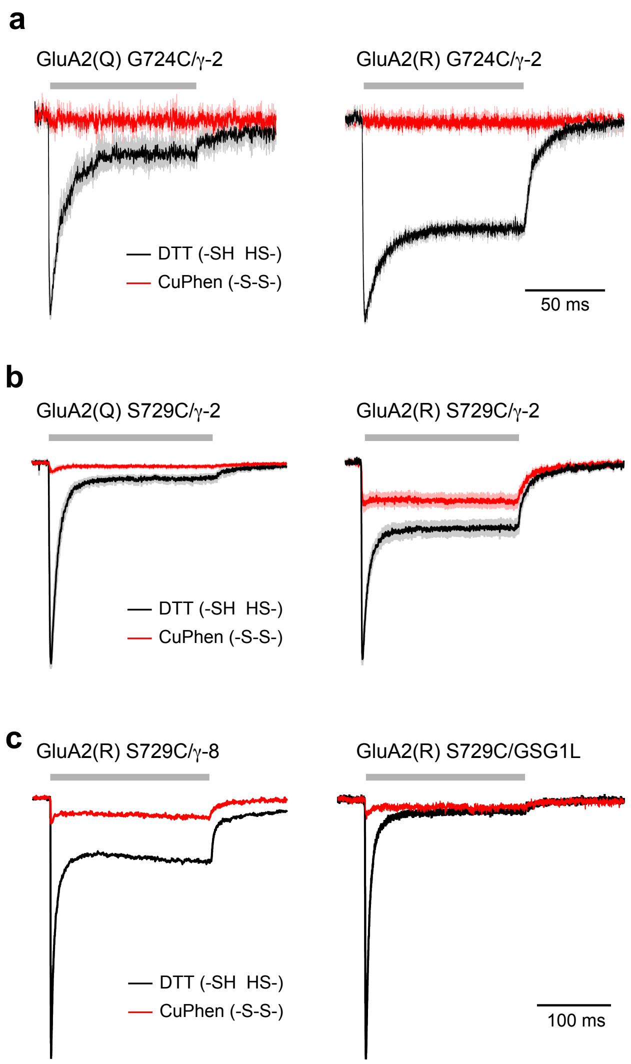

**Supplementary Figure 5. Globally averaged currents showing the effects of cross-linking for GluA2 G724C and S729C mutants**. **a**) Global averaged currents at –60 mV activated by 10 mM glutamate (100 ms; gray bar) from unedited and edited GluA2 G724C/γ-2 ($n$ = 6 and 5). Note that, GluA2(R) G724C/γ-2 is fully inhibited following cross-linking by 10 μM CuPhen. **b**) Same as a, but for GluA2 S729C/γ-2 ($n$ = 10 and 9). Note that, unlike G724C, GluA2(R) S729C is not fully inhibited following cross-linking. In each case lighter shading indicates sem. **c**) Global averaged currents showing that cross-linking of GluA2(R) S729C is unable to fully inhibit steady-state currents when the receptors are co-expressed with γ-8 or GSG1L ($n$ = 4 for both). Note that the steady-state current remaining after cross-linking (normalized to the peak current in control; $I_{\text{SS-CuPhen}}/I_{\text{Pk-DTT}}$) was 0.19, 0.07, and 0.03, for GluA2(R) S729C expressed with γ-2, γ-8 or GSG1L.

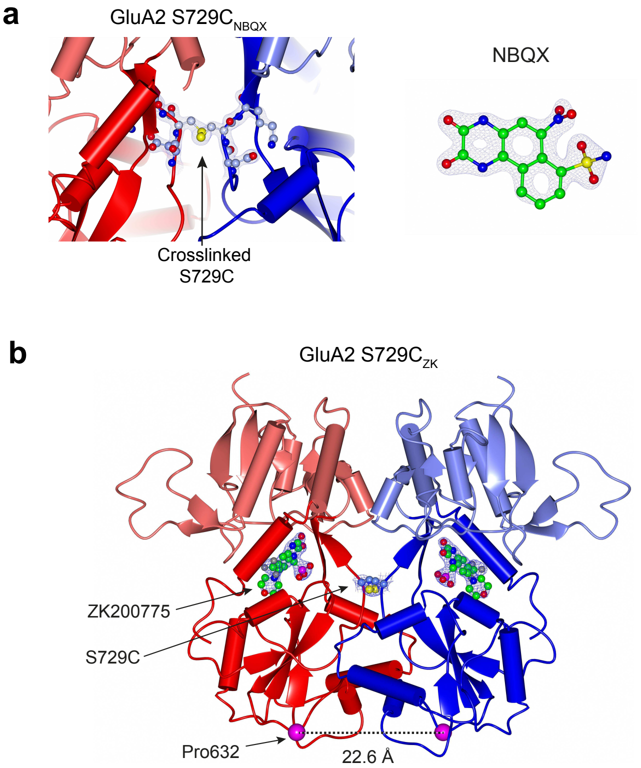

**Supplementary Figure 6. Electron densities of ligands and disulfide bonds for S729C$_{\mathbf{NBQX}}$ and S729C$_{\mathbf{ZK}}$.** **a**) $2F_o$-$F_c$ electron densities around residues 728-730, including the S729C disulfide bond (contoured at 2.0 $\sigma$) and around NBQX from S729C$_{NBQX}$ subunit 'A' (contoured at 2.5 $\sigma$). **b**) Structure of GluA2 S729C$_{ZK}$, which is highly similar to that of S729C$_{NBQX}$ (Fig. 7) (RMSD = 0.65 Å). Monomers are colored red and blue. Arrows indicate the ligand and the S729C disulfide bond (with their $2F_o$-$F_c$ electron density contoured at 2.0 $\sigma$), and the C$\alpha$ atoms of Pro632 residues (magenta spheres).

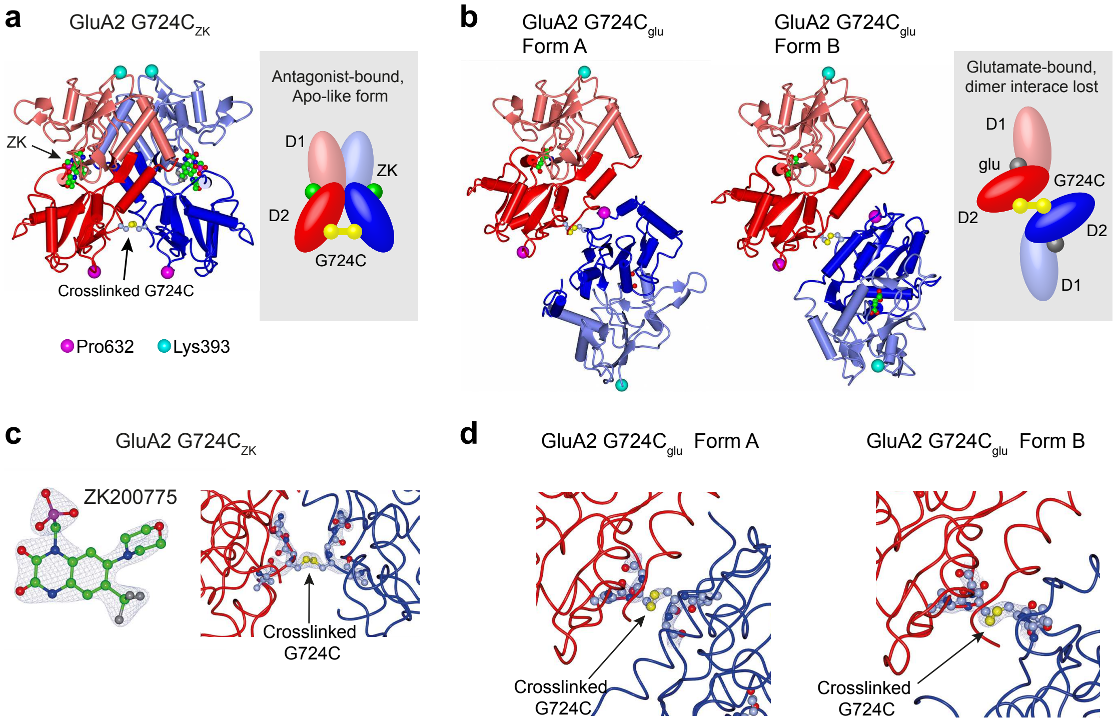

**Supplementary Figure 7. G724C cross-linking disrupts the relaxed dimer structure of LBD**. **a**) Crystal structure of the dimeric GluA2 G724C ligand binding core in the presence of ZK200775 (ZK). There were eight monomers in the unit cell, only Monomer 'A' (red) and Monomer 'H' (blue) are displayed. C$\alpha$ atoms of Lys393 positioned at the ATD-LBD linker (cyan spheres) and proline 632 at the M3-D2 linker (magenta spheres) emerge on the same sides of the dimer consistent with the expected LBD orientation within full length structures. Cartoon illustrates the position of the disulfide bond (yellow) between D2 lobes of the two monomers. **b**) Two forms of the glutamate-bound GluA2 G724C ligand binding core. D1 lobes of red monomers are displayed in the same orientation as in a. Unlike the antagonist bound form, the monomers undergo a large relative rotation compared with wild-type forms. While pairs of Pro632 residues are adjacent, Lys393 residues are positioned on opposite sides of the complex, which does not form a 'relaxed dimer' (see cartoon). **c**) $2F_o$-$F_c$ electron density around ZK200775 (subunit 'A', from a), (contoured at 2.0 $\sigma$) and the G724C disulfide bond (subunit 'A' and 'H'), (contoured at 2.5 $\sigma$). **d**) $2F_o$-$F_c$ electron density around the G724C disulfide bond (contoured at 2.0 $\sigma$) for both forms of G724C$_{glu}$.

### Supplementary Table 1

**Data collection and refinement statistics.**

| | G724C | G724C | G724C | S729C | S729C |
|---|---|---|---|---|---|
| Additives | ZK200775 | Glutamate (Form A) | Glutamate (Form B) | NBQX | ZK200775 |
| PDB code | 6FQJ | 6FQG | 6FQI | 6FQH | 6FQK |
| Data collection | | | | | |
| Beamline | DLS I24 | ESRF ID 30B | DLS I04 | DLS I04 | DLS I04 |
| Space group | P 21 21 21 | P 1 21 1 | P 41 | P 61 | P 61 |
| Cell dimensions a, b, c (Å) | 86.01, 144.29, 199.18 | 50.465, 87.655, 68.280 | 50.65, 50.65, 256.29 | 108.12, 108.12, 99.13 | 106.70, 106.70, 100.39 |
| Cell angles $\alpha$, $\beta$, $\gamma$ (Å) | 90, 90, 90 | 90, 110.223, 90 | 90, 90, 90 | 90, 90, 120 | 90, 90, 120 |
| Wavelength (Å) | 0.9686 | 0.97623 | 0.9795 | 0.9795 | 0.9795 |
| Resolution (Å) | 81.96-2.5 (2.59-2.50) | 51.73-2.34 (2.43-2.34) | 49.69-2.91 (3.01-2.91) | 28.8-1.76 (1.82-1.76) | 92.41-1.98 (2.05-1.98) |
| $R_{meas}$ | 0.121 (0.499) | 0.098 (0.769) | 0.121 (0.887) | 0.041 (1.331) | 0.06 (1.323) |
| $R_{pim}$ | 0.086 (0.353) | 0.052 (0.408) | 0.068 (0.493) | 0.016 (0.574) | 0.024 (0.509) |
| Half-set correlation $CC_{1/2}$ (%) | 0.985 (0.736) | 0.997 (0.765) | 0.995 (0.613) | 1 (0.537) | 0.997 (0.602) |
| $I/\sigma I$ | 6.8 (2.2) | 9.8 (1.8) | 10.7 (1.7) | 23.1 (1.4) | 17.2 (1.5) |
| Completeness (%) | 99.9 (100.0) | 96.9 (93.2) | 99.6 (99.9) | 100.0 (100.0) | 100.0 (99.7) |
| Multiplicity | 1.9 (1.9) | 3.4 (3.4) | 3.1 (3.1) | 6.7 (5.3) | 6.5 (6.6) |
| Refinement | | | | | |
| Resolution (Å) | 81.96-2.50 (2.59-2.50) | 51.73-2.34 (2.43-2.34) | 49.69-2.91 (3.01-2.91) | 28.8-1.76 (1.82-1.76) | 53.35-1.98 (2.05-1.98) |
| No. of reflections | 86340 | 22793 | 13987 | 64664 | 43434 |
| $R_{work}/R_{free}$ | 0.199/0.265 | 0.179/0.233 | 0.214/0.267 | 0.189/0.224 | 0.218/0.256 |
| No. of atoms | 17200 | 4166 | 4029 | 4531 | 4385 |
| Average *B*-factors – Protein | 49.04 | 50.43 | 71.43 | 50.47 | 53.80 |
| Average *B*-factors – Water | 37.49 | 46.68 | | 48.68 | 48.78 |
| Average *B*-factors – Ligand | 39.78 | 41.80 | 39.86 | 51.53 | 44.04 |
| Rmsd Bond lengths (Å) | 0.013 | 0.013 | 0.003 | 0.011 | 0.009 |
| Rmsd Bond angles (°) | 1.25 | 1.25 | 0.70 | 1.04 | 1.13 |
| Ramachandran favored/outliers (%) | 97.3/0.2 | 96.5/0.0 | 94.8/0.4 | 97.7/0.2 | 98.3/0.0 |

### Supplementary Table 2

**Details of statistical analyses.**

| Figure | Test | Description | Statistic | *P*-value | Method |
|---|---|---|---|---|---|
| **Fig. 1c** $\tau_{w, des}$ | 2-way ANOVA | Main effect of Q/R editing | $F_{1,97}$ = 111.34 | 2.20 e-16 | Welch heteroscedastic *F*-test |
| | | Main effect of auxiliary subunit type | $F_{3,97}$ = 32.30 | 1.45 e-14 | Welch heteroscedastic *F*-test |
| | | Interaction | $F_{3,97}$ = 2.84 | 0.041 | Welch heteroscedastic *F*-test |
| | Pairwise tests | GluA2(Q) *vs* GluA2(R) | $t_{15.58}$ = 10.26 | 2.48 e-08 | Welch two-sample (two-sided) t-test |
| | | GluA2(Q)/γ-2 *vs* GluA2(R)/γ-2 | $t_{43.01}$ = 2.39 | 0.021 | Welch two-sample (two-sided) *t*-test |
| | | GluA2(Q)/γ-8 *vs* GluA2(R)/γ-8 | $t_{15.00}$ = 3.43 | 0.0037 | Welch two-sample (two-sided) *t*-test |
| | | GluA2(Q)/GSG1L *vs* GluA2(R)/GSG1L | $t_{15.99}$ = 3.24 | 0.0052 | Welch two-sample (two-sided) *t*-test |
| **Fig. 1d** $I_{SS}$ | 2-way ANOVA | Main effect of Q/R editing | $F_{1,97}$ = 129.98 | 1.31 e-19 | Welch heteroscedastic *F*-test |
| | | Main effect auxiliary subunit type | $F_{3,97}$ = 58.30 | 1.24 e-21 | Welch heteroscedastic *F*-test |
| | | Interaction | $F_{3,97}$ = 58.67 | 1.02 e-21 | Welch heteroscedastic *F*-test |
| | Pairwise tests | GluA2(Q) *vs* GluA2(R) | $t_{8.95}$ = 7.70 | 3.07 e-05 | Welch two-sample (two-sided) *t*-test |
| | | GluA2(Q)/γ-2 *vs* GluA2(R)/γ-2 | $t_{33.57}$ = 14.32 | 7.35 e-16 | Welch two-sample (two-sided) *t*-test |
| | | GluA2(Q)/γ-8 *vs* GluA2(R)/γ-8 | $t_{9.70}$ = 7.76 | 1.82 e-05 | Welch two-sample (two-sided) *t*-test |
| | | GluA2(Q)/GSG1L *vs* GluA2(R)/GSG1L | $t_{8.88}$ = 1.42 | 0.19 | Welch two-sample (two-sided) *t*-test |
| **Fig. 1f** γ | Pairwise tests | GluA2(Q) *vs* GluA2(Q)/γ-2 | $t_{26.93}$ = 6.98 | 5.06 e-7 | Welch two-sample (two-sided) *t*-test |
| | | GluA2(Q) *vs* GluA2(Q)/γ-8 | $t_{6.86}$ = 3.67 | 0.016 | Welch two-sample (two-sided) *t*-test |
| | | GluA2(Q) *vs* GluA2(Q)/GSG1L | $t_{15.68}$ = 2.98 | 0.016 | Welch two-sample (two-sided) *t*-test |
| **Fig. 3a** γ | Pairwise test | GluA2(Q)/γ-2 Activation *vs* Deact. | $t_4$ = 0.51 | 0.64 | Paired *t*-test |
| **Fig. 4c** $\tau_{w, des}$ | 2-way ANOVA | Main effect of Q/R editing | $F_{1, 17}$ = 10.56 | 0.0047 | Welch heteroscedastic *F*-test |
| | | Main effect of S754D mutation | $F_{1, 17}$ = 43.19 | 4.75 e-6 | Welch heteroscedastic *F*-test |
| | | Interaction | $F_{1, 17}$ = 2.63 | 0.12 | Welch heteroscedastic *F*-test |
| | Pairwise tests | GluA2(Q)/γ-2 *vs* GluA2(Q)/γ-2 S754D | $t_{5.00}$ = 5.85 | 0.0021 | Welch two-sample (two-sided) *t*-test |
| | | GluA2(R)/γ-2 *vs* GluA2(R)/γ-2 S754D | $t_{4.00}$ = 4.64 | 0.0097 | Welch two-sample (two-sided) *t*-test |
| **Fig. 4d** $\tau_{w, recov}$ | 2-way ANOVA | Main effect of Q/R editing | $F_{1, 17}$ = 0.13 | 0.72 | Welch heteroscedastic *F*-test |
| | | Main effect of S754D mutation | $F_{1, 17}$ = 31.67 | 3.01 e-5 | Welch heteroscedastic *F*-test |
| | | Interaction | $F_{1, 17}$ = 1.65 | 0.22 | Welch heteroscedastic *F*-test |
| | Pairwise tests | GluA2(Q)/γ-2 *vs* GluA2(Q)/γ-2 S754D | $t_{6.24}$ = 4.68 | 0.0030 | Welch two-sample (two-sided) *t*-test |
| | | GluA2(R)/γ-2 *vs* GluA2(R)/γ-2 S754D | $t_{5.09}$ = 4.42 | 0.0066 | Welch two-sample (two-sided) *t*-test |
| **Fig. 4e** $I_{SS}$ | 2-way ANOVA | Main effect of Q/R editing | $F_{1, 17}$ = 65.37 | 3.16 e-7 | Welch heteroscedastic *F*-test |
| | | Main effect of S754D mutation | $F_{1, 17}$ = 28.72 | 5.56 e-5 | Welch heteroscedastic *F*-test |
| | | Interaction | $F_{1, 17}$ = 14.93 | 0.0012 | Welch heteroscedastic *F*-test |
| | Pairwise tests | GluA2(Q)/γ-2 *vs* GluA2(Q)/γ-2 S754D | $t_{5.03}$ = 4.71 | 0.0052 | Welch two-sample (two-sided) *t*-test |
| | | GluA2(R)/γ-2 *vs* GluA2(R)/γ-2 S754D | $t_{4.23}$ = 5.57 | 0.0043 | Welch two-sample (two-sided) *t*-test |
| **Fig. S1c** $\tau_{w, des}$ | Pairwise test | GluA2(Q) *vs* GluA2(Q)/γ-2 | $t_6$ = 4.44 | 0.0043 | Paired *t*-test |
| $I_{SS}$ | Pairwise test | GluA2(Q) *vs* GluA2(Q)/γ-2 | $t_6$ = 4.11 | 0.0063 | Paired *t*-test |
| **Fig. S2b** $\tau_{w, deact}$ | 2-way ANOVA | Main effect of Q/R editing | $F_{1, 37}$ = 34.70 | 8.82 e-7 | Welch heteroscedastic *F*-test |
| | | Main effect of γ-2 | $F_{1, 37}$ = 17.47 | 1.71 e-4 | Welch heteroscedastic *F*-test |
| | | Interaction | $F_{1, 37}$ = 20.64 | 5.74 e-5 | Welch heteroscedastic *F*-test |
| | Pairwise tests | GluA2(Q) *vs* GluA2(Q)/γ-2 | $t_{14.97}$ = 2.17 | 0.046 | Welch two-sample (two-sided) *t*-test |
| | | GluA2(R) *vs* GluA2(R)/γ-2 | $t_{21.72}$ = 6.06 | 4.41 e-6 | Welch two-sample (two-sided) *t*-test |
| **Fig. S3** $I_{rev}$ shift | Pairwise test | GluA2(R)/γ-2 peak *vs* steady-state | $t_6$ = 1.94 | 0.10 | Paired *t*-test |
